# EXO1-mediated DNA repair by single-strand annealing is essential for BRCA1-deficient cells

**DOI:** 10.1101/2023.02.24.529205

**Authors:** B. van de Kooij, A. Schreuder, R.S. Pavani, V. Garzero, A. Van Hoeck, M. San Martin Alonso, D. Koerse, T.J. Wendel, E. Callen, J. Boom, H. Mei, E. Cuppen, A. Nussenzweig, H. van Attikum, S.M. Noordermeer

**Affiliations:** Department of Human Genetics, Leiden University Medical Centre, Leiden, The Netherlands; Oncode Institute, Utrecht, The Netherlands; Laboratory of Genome Integrity, National Cancer Institute, National Institutes of Health, Bethesda, Maryland, USA; Centre for Molecular Medicine, University Medical Centre Utrecht, Utrecht, The Netherlands; Sequencing Analysis Support Core, Leiden University Medical Centre, The Netherlands; Hartwig Medical Foundation, Amsterdam, The Netherlands

**Keywords:** Homologous recombination, single-strand annealing, BRCA1, EXO1, synthetic lethality, DNA double-strand break repair, cancer

## Abstract

Deficiency for the repair of DNA double-strand breaks (DSBs) via homologous recombination (HR) leads to chromosomal instability and diseases such as cancer. Yet, defective HR also results in vulnerabilities that can be exploited for targeted therapy. Here, we identify such a vulnerability and show that BRCA1-deficient cells are dependent on the long-range end-resection factor EXO1 for survival. EXO1 loss results in DNA replication-induced lesions decorated by poly(ADP-ribose)-chains. In cells that lack both BRCA1 and EXO1, this is accompanied by unresolved DSBs due to impaired single-strand annealing (SSA), a DSB repair process that requires the activity of both proteins. In contrast, BRCA2-deficient cells have increased SSA, also in the absence of EXO1, and hence are not dependent on EXO1 for survival. In agreement with our mechanistic data, BRCA1-mutated tumours have elevated *EXO1* expression and contain more genomic signatures of SSA compared to BRCA1-proficient tumours. Collectively, our data indicate that EXO1 is a promising novel target for treatment of BRCA1-deficient tumours.

## Introduction

High fidelity repair of DNA double-strand breaks (DSBs) is essential in maintaining genomic integrity and preventing the onset of diseases such as cancer. Cells are equipped with several pathways to repair DSBs with varying fidelity. Homologous recombination (HR) is considered the most faithful type of repair. Loss of HR by disrupting mutations in core HR-factors such as BRCA1, BRCA2, and PALB2 is observed in many tumours of different origins and results in a distinct pattern of base substitution, mutations and rearrangements (Kandoth et al., 2013; Nik-Zainal et al., 2016). Furthermore, hereditary heterozygous mutations in these genes predispose to the development of breast, ovarian and prostate cancer (Lord and Ashworth, 2016). Besides causing tumour formation, HR deficiency also provides vulnerabilities that can be exploited for anti-cancer therapy. The reduced capacity of HR-deficient tumours to maintain genomic stability renders them sensitive to drugs that damage DNA, such as platinum compounds or PARP inhibitors (PARPi) (Lord and Ashworth, 2017). Unfortunately, cancer cells frequently acquire resistance to these types of drugs (Noordermeer and van Attikum, 2019). This indicates a necessity to develop additional single-drug or combinatorial therapies to reduce the chance of relapse.

To enable repair of DSBs by HR, the flanking DNA needs to be resected to expose stretches of single stranded DNA (ssDNA). The ssDNA is then bound by the core HR-factor RAD51 which facilitates the search for an intact homologous sequence, most frequently the sister chromatid, that is subsequently invaded and functions as a template for repair. BRCA1 has been shown to play a role at multiple steps during HR. It facilitates RAD51 loading onto the resected DNA by recruitment of PALB2-BRCA2 and direct interaction with RAD51 (Scully et al., 2019; Zhao et al., 2017). In addition, BRCA1 stimulates the upstream process of DNA end-resection (Callen et al., 2020; Chandramouly et al., 2013; Cruz-Garcia et al., 2014).

Resection is initiated by the short-range end-resection factors MRE11 and CTIP which remove up to 300 nucleotides of ssDNA (Garcia et al., 2011). Long-range end-resection subsequently can process thousands of base pairs (bps) and is mediated either via the exonuclease EXO1, or the flap endonuclease DNA2, which cooperates with the helicase BLM (Gnugge and Symington, 2021). Interestingly, while short-range end-resection has been shown to be essential for HR, the role of long-range end-resection is more enigmatic (Garcia et al., 2011; Lu et al., 2016; Symington, 2016). Several studies showed that long-range end-resection plays a stimulatory role in HR and checkpoint activation (Karanja et al., 2012; Kimble M. T., 2022; Misenko et al., 2018). In contrast, others have shown that HR is not affected by loss of long-range end-resection proteins (Chen et al., 2017; Ochs et al., 2016; Patel et al., 2017; Tomimatsu et al., 2017; van de Kooij et al., 2022).

In addition to HR, two other DSB repair pathways are also dependent on end-resection: alternative end-joining (alt-EJ) and single-strand annealing (SSA). During alt-EJ, limited end-resection exposes regions of microhomology flanking the DSB-ends which anneal during repair, generally resulting in deletion of the intermittent sequence (Kelso et al., 2019; Mateos-Gomez et al., 2015; van Schendel et al., 2021; Yousefzadeh et al., 2014). SSA requires more extensive long-range end-resection to expose larger stretches of (imperfect) homology which can be located at distal regions up- and downstream of the DSB. The resected DNA is bound by the mediator protein RAD52 that drives annealing of the homologous regions, followed by the removal of the non-homologous intermittent ssDNA and subsequent repair by polymerases and ligases. Compared to alt-EJ, DSB repair by SSA generally leads to larger deletions (Scully et al., 2019).

Although the contribution of SSA and alt-EJ to the repair of physiological DSBs in healthy cells is not fully understood, loss of either the alt-EJ factor POLQ or the SSA factor RAD52 is synthetically lethal (SL) with BRCA1 or BRCA2 loss. This implies that DSB repair by alt-EJ or SSA is essential for survival of HR-deficient cells (Ceccaldi et al., 2015; Feng et al., 2011; Lok et al., 2013; Mateos-Gomez et al., 2015). However, recent studies suggested that increased ssDNA gap formation in absence of PolQ may underlie the synthetic lethality with BRCA-deficiency (Belan et al., 2022; Mann et al., 2022; Schrempf et al., 2022). BRCA-deficient cells are prone to accumulate ssDNA gaps, and toxic levels of these lesion are associated with sensitivity to chemotherapy and PARPi (Cong et al., 2021; Paes Dias et al., 2021; Panzarino et al., 2021). Similarly, for RAD52, the synthetic lethal interaction with BRCA deficiency might not be driven by loss of SSA, but rather by a potential function of RAD52 in BRCA-independent HR (Feng et al., 2011; Lok et al., 2013), questioning the importance of SSA for survival of BRCA-deficient cells.

In our search for novel targets to treat BRCA-deficient tumours, we studied the role of resection factors in BRCA-deficient cells. We show that long-range end-resection mediated by EXO1 is essential for BRCA1-deficient cells, but not BRCA2-deficient cells. Our mechanistic data indicate that EXO1 depletion induces DNA lesions in both genetic backgrounds, yet only BRCA1-deficient cells cannot tolerate EXO1 loss due to an inability to overcome this damage. Thus, we uncover a novel vulnerability of BRCA1-deficient tumour cells and we propose EXO1 as a novel therapeutical target for BRCA1-deficient tumours.

## Results

### Long-range end-resection factors are essential for BRCA1-deficient cells

A CRISPR screen that we performed previously, designed to identify essential genes in isogenic BRCA1-deficient versus BRCA1-proficient RPE1 cell lines (Adam et al., 2021) allowed us to study the genetic interaction between BRCA1 deficiency and the loss of resection factors. As indicated by the high CCA score, the long-range end-resection factors EXO1 and BLM were essential in BRCA1-deficient cells, but not in BRCA1-proficient cells (Figure 1A), to a similar extent as previously described SL interactors such as APEX2 and PARP1 (Figure 1A) (Alvarez-Quilon et al., 2020; Bryant et al., 2005; Farmer et al., 2005). DNA2, another long-range end-resection factor, was essential in both BRCA1-deficient and -proficient cells, in accordance with previous findings (Appanah et al., 2020; Hart et al., 2017; Lin et al., 2013), explaining the low CCA score (Figure 1A). In contrast to these long-range end-resection factors, the short-range end-resection factors MRE11 and CTIP did not show significant SL interactions with BRCA1 deficiency (Figure 1A).

**Figure 1.**
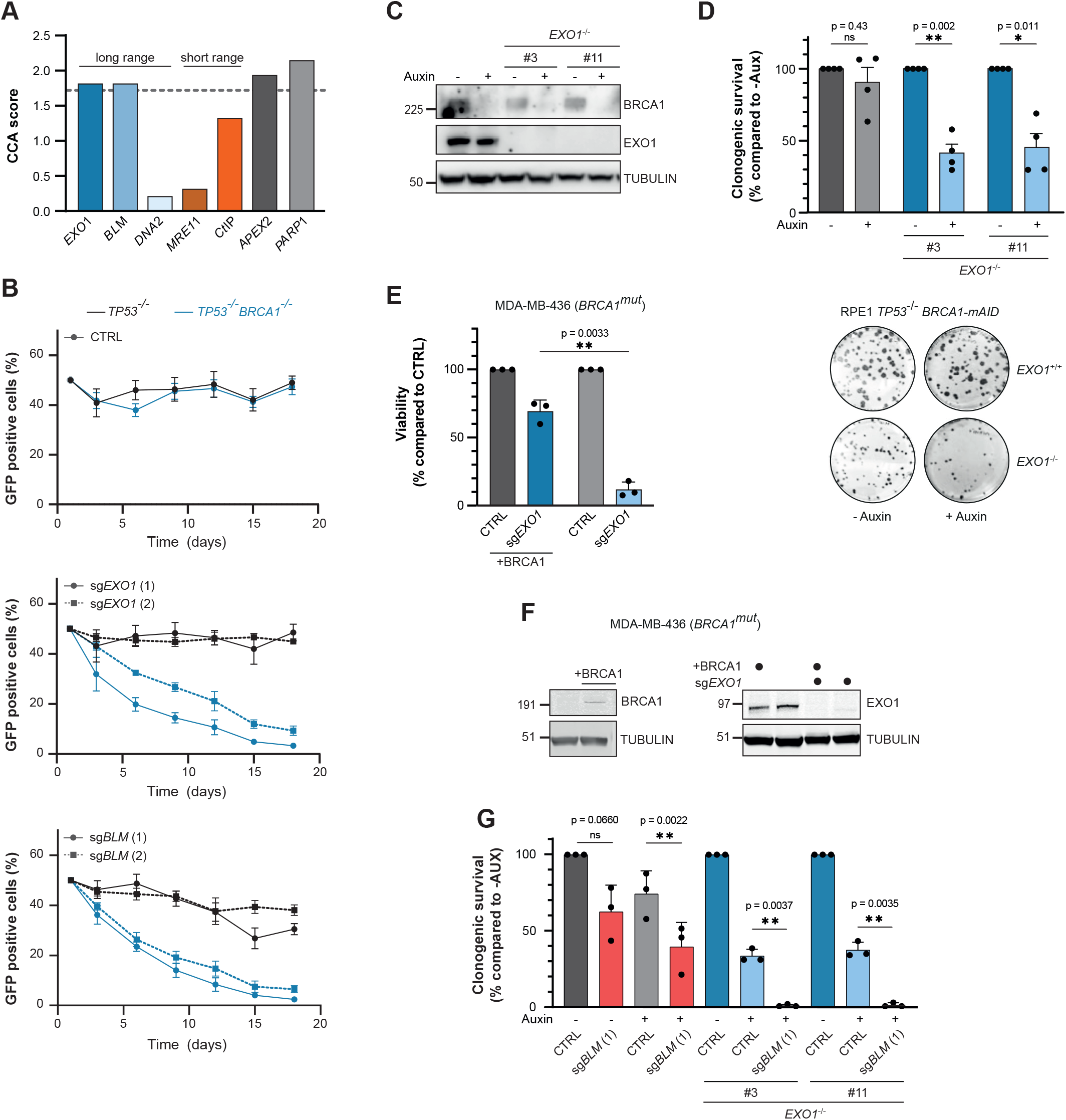
EXO1 loss is synthetically lethal with BRCA1-deficiency. **(A)** Selected results of a gene essentiality screen in BRCA1-proficient and -deficient RPE1 cells (Adam *et al.*, 2021). Plotted is the CCA score: a higher score indicates a unique essentiality in BRCA1-deficient cells compared to -proficient cells. Dashed line indicates the cut-off for a significant CCA score (based on Adam *et al.*, 2021). **(B)** RPE1 hTERT cells expressing Cas9, either *TP53*^-/-^ (black lines) or *TP53*^-/-^ *BRCA1*^-/-^ (blue lines), were infected with indicated sgRNA together with GFP, or with an empty vector together with mCherry. GFP- and mCherry-positive cells were mixed 1:1, and the frequency of GFP-positive cells in the population was determined at multiple time points (n=4, mean±SEM). Western blot of lysates shown in supplemental figure 1A. **(C)** RPE1 hTERT *PAC*^-/-^ *TP53*^-/-^ cells were genetically modified to generate a *BRCA1-mAID-GFP* fusion gene at the endogenous *BRCA1*-locus. In this genetic background, two clonal *EXO1*^-/-^ cell lines were generated using gene editing techniques. Cells were treated with auxin (500 μM) for 48 hours or left untreated, and lysates were analysed by western blotting. **(D)** The cell lines described in figure 1C were treated with 500 μM auxin, or left untreated, and clonogenic survival was determined. Lower panel shows a representative experiment, top panel shows the quantification (n=4, mean+SEM, *p<0.05, **p<0.01, paired t-test). **(E)** *BRCA1*-mutated MDA-MB-436 cells, either WT or reconstituted with BRCA1, were infected with empty vector (CTRL) or *EXO1*-targeting sgRNA and viability was measured using CellTiter-Glo (n=3, mean+SD, **p<0.01, paired t-test). **(F)** Lysates of the MDA-MB-436 cell lines described in figure 1E were analysed by western blotting. **(G)** RPE1 hTERT *TP53*^-/-^ *BRCA1-mAID-GFP* cell lines were virally transduced to express *Cas9* cDNA and the indicated gRNAs, followed by a clonogenic survival assay in presence or absence of 500 μM Auxin (n=3, mean+SD, **p<0.01, paired t-test). Western blot of lysates shown in supplemental figure 1L.

To validate these results, we performed a competitive growth assay as described before (Noordermeer et al., 2018). In short, GFP-positive cells, depleted for the gene-of-interest, were mixed 1 to 1 with mCherry-positive non-depleted cells, and the composition of the mixed population was monitored over time. Confirming data from the CRISPR screen, depletion of EXO1 or BLM by CRISPR/Cas9 strongly reduced the proliferation of BRCA1-deficient cells but not of BRCA1-proficient cells (Figure 1B, Supplemental figure 1A). CTIP loss was toxic to both BRCA1-deficient and -proficient cells, although BRCA1-deficient cells seemed more sensitive to CTIP loss as was described previously (Przetocka et al., 2018). Loss of MRE11 reduced proliferation of both BRCA1-proficient and -deficient cells (Supplemental figure 1B,C). Together, these data indicate that short-range resection is essential in all contexts, which is in agreement with previous reports (Buis et al., 2008; Polato et al., 2014), while long-range end-resection factors are essential specifically in a BRCA1-deficient background.

To enable mechanistic studies on the synthetic lethal interaction between BRCA1-deficiency and loss of long-range end-resection, we generated an RPE1 hTERT cell line containing an auxin-inducible degron (AID) fused to the *BRCA1* gene. In this background, we generated clonal *EXO1* knock-out cell lines using CRISPR/Cas9 (Figure 1C). Confirming our previous data, clonogenic survival of *EXO1^-/-^* cells was strongly reduced upon auxin-induced BRCA1 depletion while the genetic depletion of EXO1 by itself only mildly affected cellular proliferation (Figure 1D). Importantly, complementation of the *EXO1^-/-^* cells with full-length EXO1 cDNA rescued the toxicity of auxin-induced BRCA1 depletion (Supplemental figure 1D,E). Moreover, complementation of *BRCA1* knock-out cells with full length BRCA1 restored the viability upon EXO1 depletion (Supplemental figure 1F,G), demonstrating that the synthetic lethality is not due to off-target effects of the genetic approaches to deplete EXO1 and BRCA1. Finally, we confirmed the SL interaction in the patient-derived *BRCA1*-mutated MDA-MB-436 cells (Figure 1E-F, Supplemental figure 1H,I) (Elstrodt et al., 2006; Johnson et al., 2013) and in BRCA1-deficient (BRCA1 Δ exon 11) (Callen et al., 2020) mouse embryonic fibroblasts (Supplemental figure 1J,K), showing that the interaction is not cell type-specific.

Similar to EXO1 loss, clonogenic survival was impaired for cells depleted of both BLM and BRCA1 using our *BRCA1-mAID* cell line (Figure 1G, Supplemental figure 1L). Interestingly, BLM-depletion even further reduced clonogenic survival of BRCA1-depleted *EXO1^-/-^* cells (Figure 1G). These results indicate that BLM and EXO1 have unique, additive functions in promoting survival of BRCA1-deficient cells. BLM mutations cause the autosomal recessive Bloom syndrome which is linked with growth deficiency and cancer predisposition (German, 1993), and BLM deficiency is embryonically lethal in mice (Chester et al., 1998). Although EXO1 deficiency leads to phenotypes such as sterility and lymphomas, these mice are born at normal mendelian ratio (Rein et al., 2015; Wang et al., 2022; Wei et al., 2003). These data, combined with the observed growth reduction upon BLM loss in BRCA1-proficient cells (Figure 1B,G), suggests that targeting EXO1 might be favoured over targeting BLM as an approach to treat BRCA1-deficient tumours. Hence, we decided to focus on EXO1 in this study.

### BRCA1-mutated tumours exhibit elevated EXO1 expression

The dependency of BRCA1-deficient tumour cells on SL interactors frequently manifests itself by high levels of expression or activity of the interactor, as has been shown for POLQ and PARP1 (Ceccaldi et al., 2015; Gottipati et al., 2010). To investigate whether such correlation exists between EXO1 and BRCA1 deficiency, *EXO1* expression levels and *BRCA1* mutation status were extracted from two previously published breast cancer cohorts (Nik-Zainal et al., 2016; Staaf et al., 2019). Indeed, in both the triple-negative breast cancer cohort and the pan-breast cancer cohort, *BRCA1*-mutated or *BRCA1* promoter-hypermethylated tumours showed elevated *EXO1* expression compared to *BRCA1*-wildtype tumours (Figure 2A,B). *BLM* also showed increased expression in BRCA1-deficient tumours and the increased expression of the two genes was comparable to the increase in *POLQ* expression. *MRE11* expression was not increased in BRCA1-deficient breast tumour samples, consistent with the lack of an SL interaction with BRCA1 (Figure 2A,B).

**Figure 2.**
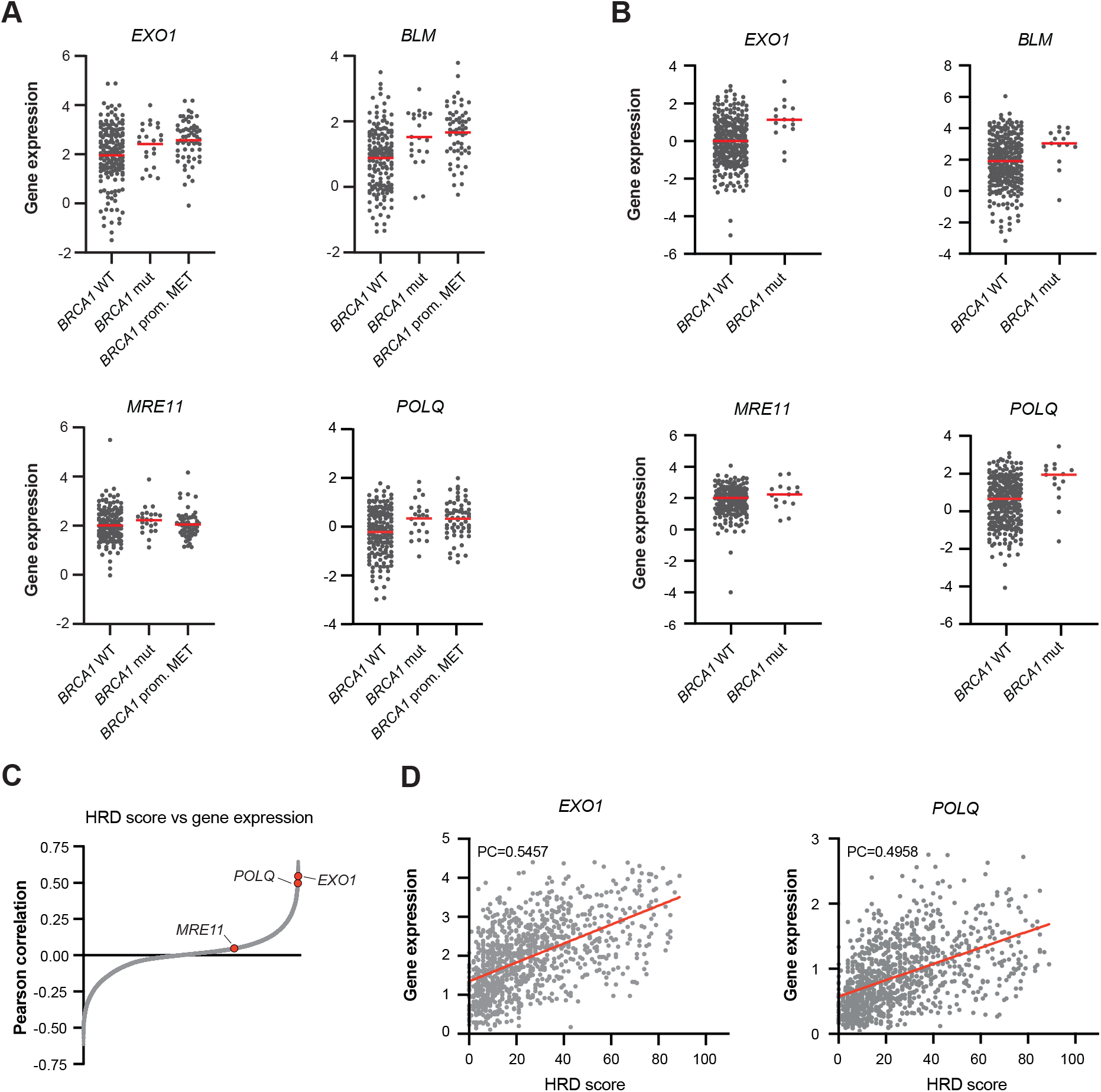
Elevated EXO1 expression in BRCA1-deficient tumours. **(A)** Primary breast tumour samples (Staaf *et al.*2019, n=247) were divided in groups based on the presence of biallelic *BRCA1* mutations (*BRCA1* mut) or *BRCA1* promotor hypermethylation (*BRCA1* prom. MET), and gene expression of indicated genes were plotted. **(B)** As in figure 2A, but for another cohort of breast cancer patients in which groups were divided on the presence of deleterious *BRCA1* mutations (Nik-Zainal *et al.*, 2016, n=342). **(C)** For tumour samples of the TCGA breast cancer dataset (n=1,048), the correlation between the HRD score and expression level of an individual gene was determined. Plotted are the Pearson correlation coefficient for each of 60,000 measured transcripts. **(D)** HRD score and expression of the indicated genes were plotted for each tumour sample in the TCGA breast cancer dataset (n=1,048). Red line is the linear regression curve, PC=Pearson correlation coefficient.

To extrapolate the elevated *EXO1* expression found in *BRCA1*-mutated tumours to HR-deficient tumours more generally, the TCGA breast cancer dataset was used to correlate genome-wide gene expression to homologous recombination deficiency (HRD) scores (Knijnenburg et al., 2018). A strong positive correlation was observed between *EXO1* expression and HRD score, while such a correlation was absent for *MRE11* (Figure 2C). In fact, *EXO1* was among the top hits that showed the strongest positive correlation with HRD score with a Pearson correlation even higher than for *POLQ* (Figure 2D). These results indicate that elevated *EXO1* expression correlates with HR deficiency beyond BRCA1-mutations, implying other HR-deficient cells might also depend on EXO1 function.

### EXO1 loss is not synthetically lethal in BRCA2-deficient cells

To explore whether other causes of HR deficiency result in dependency on EXO1 activity, we again examined the results of the CRISPR gene essentiality screens from Adam *et al.* which included BRCA2-deficient cells as well (Adam et al., 2021). Surprisingly, no SL interaction was identified between BRCA2 deficiency and EXO1 loss (Figure 3A). This is consistent with another recently published CRISPR screen in BRCA2-depleted HEK293A cells that also failed to identify a dependency on EXO1 expression (Tang, 2021). To validate these results, we depleted EXO1 in WT or *BRCA*2^-/-^ DLD1 cells using CRISPR/Cas9. In line with the screens, EXO1 depletion did not significantly decrease the viability of BRCA2-deficient cells compared to WT (Figure 3B,C). As a second approach, we depleted EXO1 in H1299 cells containing a doxycycline (dox) inducible *BRCA2* shRNA (Tacconi et al., 2017). Similarly, EXO1 loss did not affect the clonogenic survival of BRCA2-depleted cells compared to control cells (Figure 3D,E). Together, our results show that loss of EXO1 is toxic to BRCA1-deficient cells, but not to BRCA2-deficient cells.

**Figure 3.**
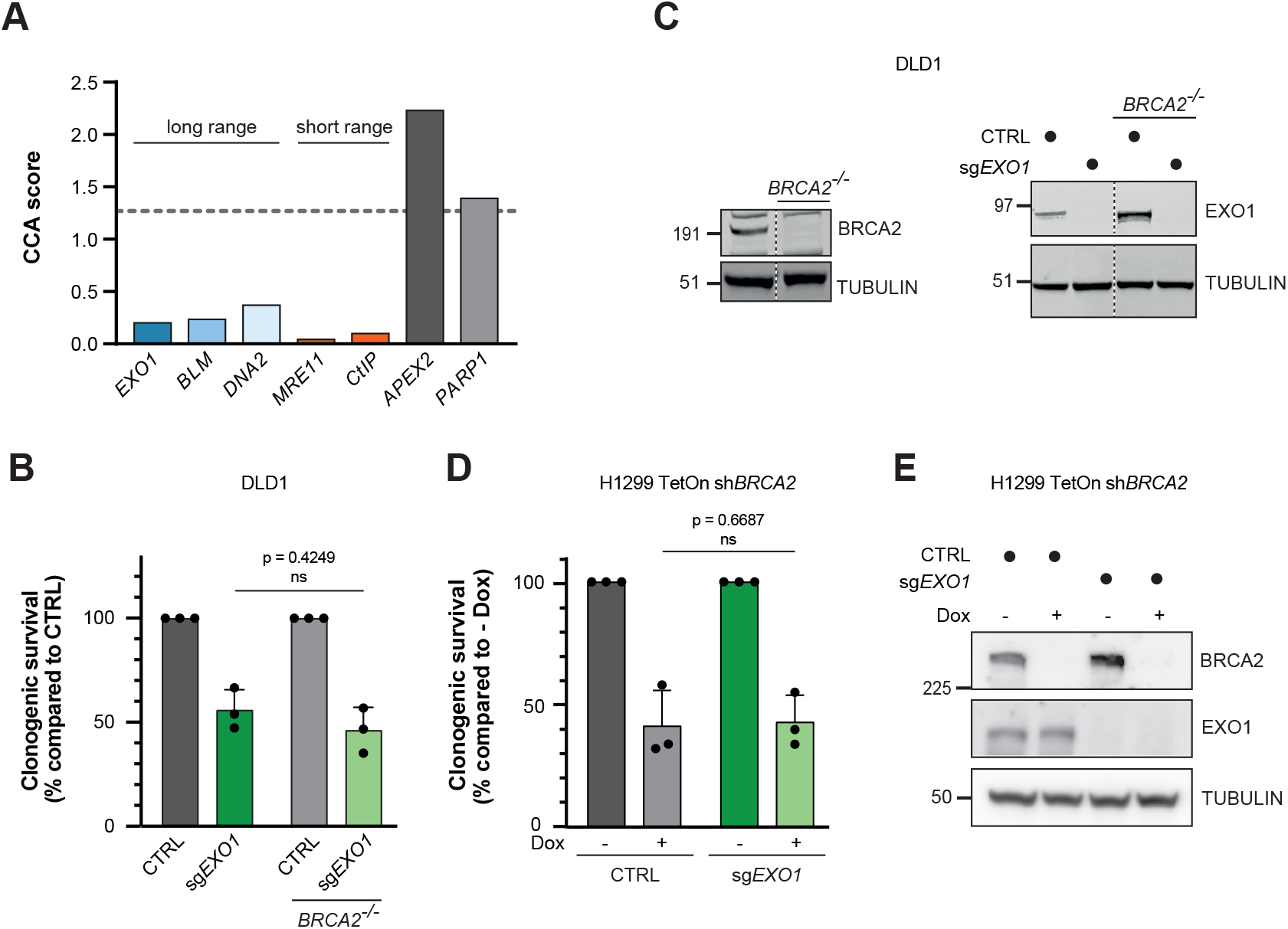
EXO1 loss is not lethal in BRCA2-deficient cells. **(A)** A published gene essentiality screen in BRCA2-proficient and -deficient RPE1 cells (Adam *et al.*, 2021) was mined to extract the CCA scores for the indicated genes. A higher CCA score indicates a unique essentiality in BRCA2-deficient cells compared to proficient cells. Dashed line indicates the cut-off for a significant CCA score (based on Adam *et al.*, 2021). **(B)** DLD1 WT and *BRCA2*^-/-^ cells were infected with empty vector (CTRL) or *EXO1*-targeting sgRNA and viability was measured using CellTiter-Glo (n=3, mean+SD, paired t-test). **(C)** Lysates of the indicated DLD1 cells indicated in figure 3B analysed by western blotting. Samples were run on the same gel, dashed line indicates removal of non-relevant lanes post-imaging. **(D)** H1299 cells carrying a doxycycline-inducible *BRCA2* shRNA were transduced with an AAVS1-targeting (CTRL) or *EXO1*-targeting sgRNA, followed by a clonogenic survival assay in absence or presence of 10 μg/mL doxycycline (n=3, mean+SD, paired t-test). **(E)** Western blot analysis of the lysates of the H1299 TetOn sh*BRCA2* cells studied in figure 3D.

### Genomic instability in BRCA1-deficient cells is exacerbated by EXO1 depletion

To elucidate the mechanism behind the synthetic lethality between BRCA1 and EXO1 loss, we investigated the effect of depleting BRCA1, EXO1, or both on genomic stability. For this, we studied spontaneous formation of micronuclei and chromosomal aberrations in our *BRCA1-mAID* cell lines with and without EXO1. In the BRCA1-proficient background, EXO1 loss did not affect micronuclei levels (Figure 4A). Auxin-induced depletion of BRCA1 increased micronuclei formation in the EXO1-proficient cells, but even more so in the EXO1-deficient cells (Figure 4A). This effect was phenocopied by genetic EXO1 depletion in a clonal *BRCA1* knock-out cell line (Figure 4B). Additionally, *BRCA1-mAID* cells co-depleted for EXO1 and BRCA1 showed an elevated number of chromosomal aberrations compared to cells depleted for either factor alone (Figure 4C). The increase was predominantly caused by chromatid aberrations, rather than chromosome aberrations (Figure 4D), suggesting these cells suffer from S-phase induced DNA damage.

**Figure 4.**
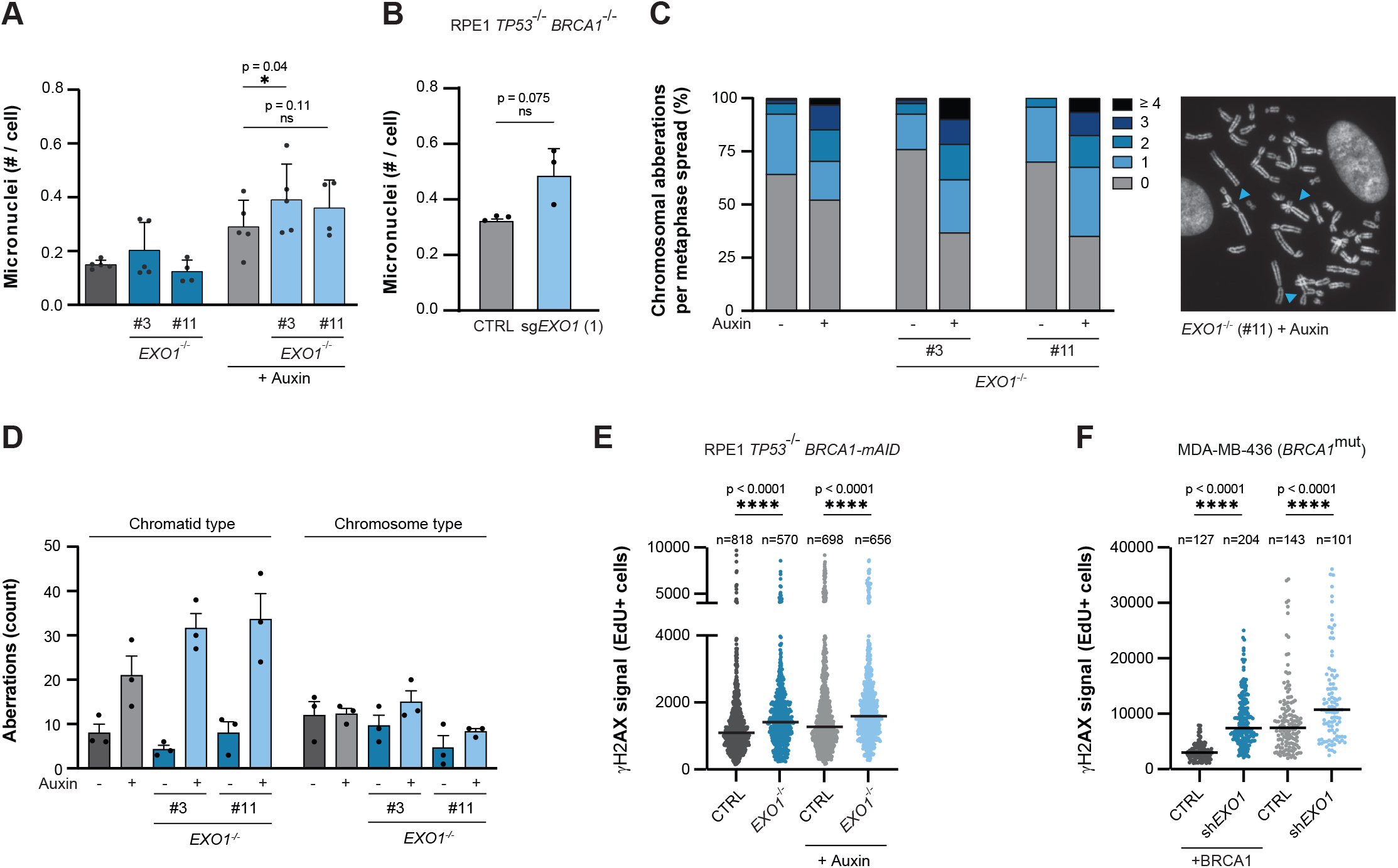
Genomic instability in BRCA1-deficient cells is exacerbated by EXO1 depletion. **(A)** RPE1 hTERT *TP53*^-/-^ *BRCA1-mAID-GFP* cells, either *EXO1*^+/+^ or *EXO1*^-/-^, were treated with 500 μM auxin for 48 hours to deplete BRCA1 or left untreated. This was followed by Hoechst staining and microscopic quantification of the number of micronuclei (n=5, mean+SD, *p<0.05, ratio paired t-test). **(B)** RPE1 hTERT *TP53*^-/-^ *BRCA1*^-/-^ cells expressing Cas9 were infected with *sgEXO1*. Next, DAPI-stained nuclei were analysed by microscopy to quantify the number of micronuclei (n=3, mean+SD, ratio paired t-test). **(C)** RPE1 hTERT *TP53*^-/-^ *BRCA1-mAID-GFP*, either *EXO1*^+/+^ or *EXO1*^-/-^, were treated with 500 μM auxin for 48 hours to deplete BRCA1 or left untreated. Next, metaphase spreads were analysed for chromosomal aberrations. Right panel shows a representative experiment with blue arrows indicating aberrations, left panel shows the quantification (n=3, >40 metaphases per replicate, mean). **(D)** Data depicted in panel C was re-plotted to show the absolute number of chromosomal aberrations, either chromatid type or chromosome type. **(E)** RPE1 hTERT *TP53*^-/-^ *BRCA1-mAID-GFP* cells, either *EXO1*^+/+^ or *EXO1*^-/-^, were treated with 500 μM auxin for 48 hours to deplete BRCA1 or left untreated. Nuclear γH2AX intensity in S-phase (EdU^+^) cells was analysed by IF microscopy. A representative of two independent experiments is shown, black line indicates median (****p<0.0001, Mann-Whitney). **(F)** Nuclear γH2AX intensity in S-phase (EdU^+^) cells analysed by IF microscopy in *BRCA1*-mutated MDA-MB-436 cells, either WT or reconstituted with BRCA1 cDNA, infected with empty vector (CTRL) or EXO1-targeting shRNA. A representative of two independent experiments is shown, black line indicates median (****p<0.0001, Mann-Whitney).

To study whether the chromosomal instability correlates with DNA damage, we measured γH2AX levels in the different genomic backgrounds under normal growth conditions. In both RPE1 and MDA-MB-436 cells, BRCA1 deficiency or EXO1 loss increased γH2AX levels in S-phase cells (Figure 4E,F). Importantly, the increase was exacerbated in cells lacking both BRCA1 and EXO1 (Figure 4E,F), suggesting that these cells suffer from unresolved DNA damage.

### EXO1^-/-^ cells show increased PARylation

To pinpoint which function of BRCA1 is essential for survival in the absence of EXO1, we assessed cell viability in cells co-depleted of 53BP1. Loss of 53BP1 restores HR (Bouwman et al., 2010; Bunting et al., 2010) and prevents ssDNA gap formation in *BRCA1^-/-^* cells (Paes Dias et al., 2021), while it does not rescue the destabilization of replication forks due to *BRCA1^-/-^* deficiency (Ray Chaudhuri et al., 2016). Interestingly, loss of 53BP1 fully restored clonogenic survival of EXO1-depleted BRCA1 KO cells (Figure 5A, Supplemental figure 2A). Hence, these experiments imply two features of BRCA1 loss that might induce toxicity in EXO1-depleted cells: defective HR or an increase in ssDNA gaps.

**Figure 5.**
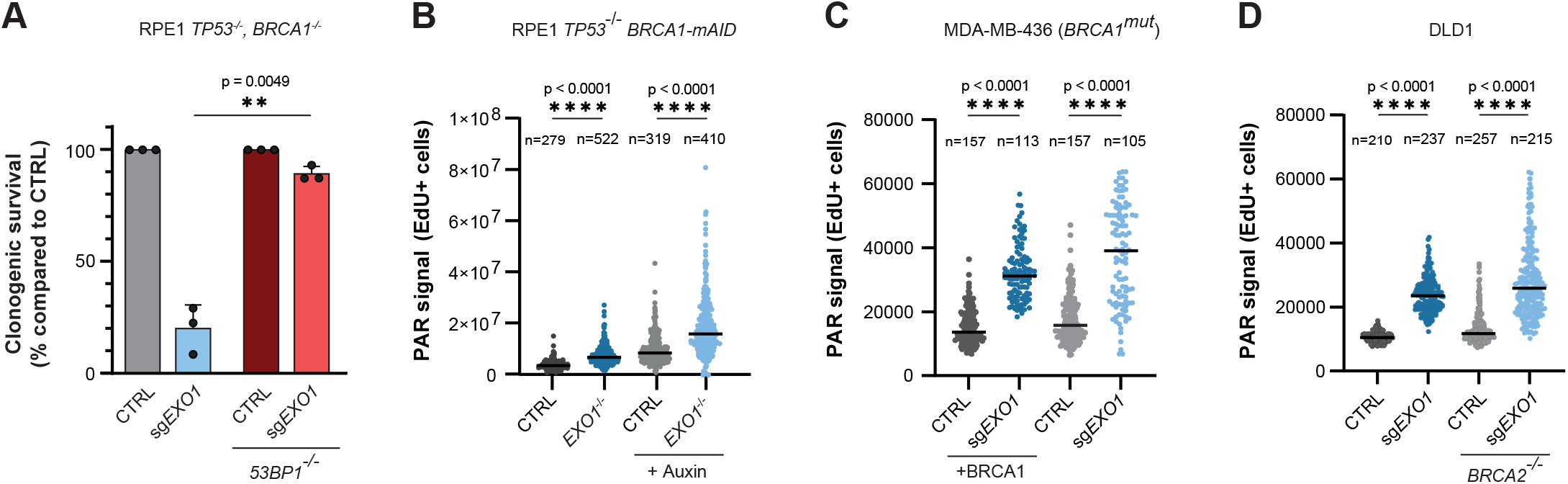
EXO1-depletion causes S-phase PAR-signalling. **(A)** RPE1 hTERT *TP53*^-/-^ *BRCA1*^-/-^ cells expressing Cas9, either *TP53BP1*^+/+^ or *TP53BP1*^-/-^, were transduced with an AAVS1-targeting (CTRL) or *EXO1*-targeting sgRNA, followed by a clonogenic survival assay (n=3, mean+SD, **p<0.01, paired t-test). Western blot of lysates shown in supplemental figure 2A. **(B)** Indicated RPE1 cell lines were incubated with EdU and PARGi for 30 minutes, followed by IF microscopy to analyse PAR formation in EdU-positive cells. A representative of two independent experiments is shown, black line indicates median (****p<0.0001, Mann-Whitney). **(C)** As in figure 5B, but for *BRCA1*-mutated MDA-MB-436 cells, either WT or reconstituted with *BRCA1* cDNA, infected with empty vector (CTRL) or *EXO1*-targeting sgRNA. A representative of three independent experiments is shown, black line indicates median (****p<0.0001, Mann-Whitney). **(D)** As in figure 5B, but for DLD1 cells, either WT or *BRCA2*^-/-^. A representative of three independent experiments is shown, black line indicates median (****p<0.0001, Mann-Whitney).

First, we examined ssDNA gap formation in cells depleted for BRCA1 and/or EXO1. To investigate this, we measured poly(ADP-ribosyl)ation (PARylation), a post-translational modification catalysed by the poly(ADP-ribose) polymerase (PARP) enzymes at sites of ssDNA breaks and gaps (Eustermann et al., 2015; Hanzlikova and Caldecott, 2019). Interestingly, EXO1 loss led to a substantial increase in S-phase PAR intensity in BRCA1-proficient cells which was even stronger in BRCA1-deficient cells (Figure 5B,C). The PAR signal co-localised with nascent DNA labelled by EdU (Supplemental figure 2B), suggesting that EXO1-loss induces replication-associated gaps in the lagging strand that trigger PARP activity (Hanzlikova et al., 2018; Vaitsiankova et al., 2022). Of note, nuclear intensity of incorporated BrdU detected under native conditions, was similar in all genetic backgrounds (Supplemental figure 2C). This likely indicates that the PAR-decorated lesions are nicks or small gaps below the detection threshold of the BrdU assay. Unexpectedly, EXO1 loss induced a similar increase in the S-phase PAR levels in *BRCA2^-/-^* cells (Figure 5D). Together, these results suggest that EXO1 loss generates DNA lesions in S-phase that can be tolerated in BRCA1-proficient and BRCA2-deficient cells, but not in BRCA1-deficient cells.

### The role of EXO1 in SSA is essential for survival in BRCA1-deficient cells

We next focused on determining which cellular function of EXO1 is required for survival of BRCA1-deficient cells. Besides its function in long-range end-resection during DSB repair, EXO1 also plays a role in mismatch repair (MMR). However, the loss of other core MMR factors (MLH1, MSH2, MSH3, MSH6, PMS1, PMS2) did not result in lethality in BRCA1-deficient cells in the aforementioned CRISPR screen (Adam et al., 2021) (Figure 6A). This therefore argues against EXO1’s role in MMR being required for the survival of BRCA1-deficient cells.

**Figure 6.**
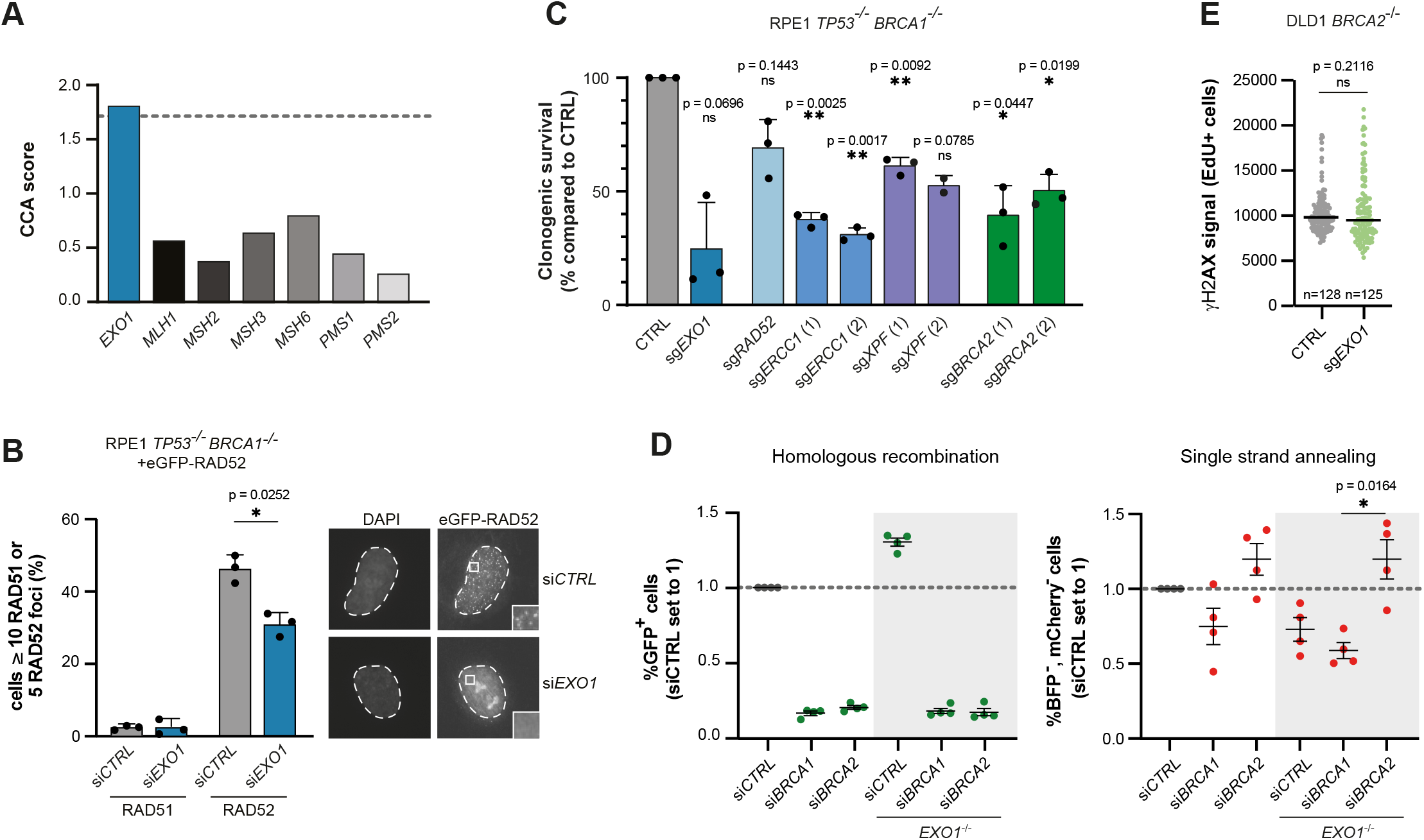
Survival of BRCA1-deficient cells is dependent on the function of EXO1 in SSA. **(A)** Results of a gene essentiality screen in BRCA1-proficient and -deficient cells (Adam *et al.*, 2021) were mined to extract the CCA scores for the indicated genes. A higher CCA score indicates a unique essentiality in BRCA1-deficient cells compared to proficient cells. Dashed line indicates the cut-off for a significant CCA score (based on Adam *et al.*, 2021). **(B)** Indicated RPE1 cells were transfected with a control (siCTRL) or *EXO1*-targeting siRNA, followed by treatment with IR and subsequent IF microscopy to analyse either RAD51 or RAD52 foci formation. Left panel shows quantification (n=3, mean+SD, *p<0.05, paired t-test) and right panel shows representative microscopy images of eGFP-RAD52 foci. Western blot of lysates shown in supplemental figure 3A. **(C)** Clonogenic survival assay of Cas9-expressing RPE1 hTERT *TP53*^-/-^ *BRCA1*^-/-^ cells that were transduced to express the indicated sgRNAs (n=3, mean+SD, *p<0.05, **p<0.01, one-way ANOVA, post-hoc Dunnett’s, compared to CTRL). Western blot of lysates shown in supplemental figure 3D. **(D)** HEK 293T cells carrying the DSB-Spectrum_V3 reporter, either WT or *EXO1*^-/-^, were transfected with indicated siRNAs, followed by a second round of transfection with a *Cas9* cDNA and *BFP* sgRNA targeting the reporter locus. Next, cells were analysed by flow cytometry to quantify repair by the indicated pathways (n=4, mean±SEM, *p<0.05, ratio paired t-test). Western blot of lysates shown in supplemental figure 3E. **(E)** Nuclear γH2AX intensity in S-phase (EdU^+^) cells analysed by IF microscopy in *BRCA2*^-/-^ cells infected with empty vector or *sgEXO1*. A representative of two independent experiments is shown, black line indicates median.

We thereafter considered that BRCA1-deficient cells depend on the function of EXO1 in long-range end-resection during DSB repair. Using our newly developed multi-pathway reporter for DSB repair, we recently found that long-range end-resection mediated by EXO1 is not required for HR but promotes SSA (van de Kooij et al., 2022). To substantiate these findings in a BRCA1-deficient setting, we examined ionising irradiation-induced foci (IRIF) of RAD51 and RAD52 in our BRCA1-deficient cell lines as a read-out of HR and SSA, respectively. As expected, RAD51 IRIF were absent in BRCA1-deficient cells (Figure 6B). Depletion of EXO1 using an siRNA did not abrogate RAD51 IRIF further, whereas it decreased the number of RAD52-GFP IRIF in these cells (Figure 6B, Supplemental figure 3A). This decrease in RAD52-GFP IRIF correlated with a decrease in end-resection caused by EXO1 loss, as demonstrated by reduced levels of RPA phosphorylation (S4/S8) (Supplemental figure 3B). These data indicate that the cells deficient for both BRCA1 and EXO1 have a reduced capacity to repair DSBs due to a defect in both HR and SSA.

The decrease of RAD52 IRIF after EXO1 loss prompted us to investigate whether BRCA1-deficient cells might rely on EXO1-mediated SSA for survival. To validate the importance of SSA in these cells, we looked at the toxicity upon loss of SSA factors other than EXO1. To this end, we used CRISPR/Cas9 to deplete the SSA-factors RAD52, ERCC1, and XPF (Al-Zain and Symington, 2021) in BRCA1-deficient RPE1 cells and measured their clonogenic survival capacity. Similar to the loss of EXO1, loss of all other tested SSA factors induced lethality in *BRCA1*^-/-^ cells (Figure 6C, Supplemental figure 3D). The toxicity of RAD52 loss has been ascribed to a function for RAD52 in promoting residual HR in BRCA1-deficient cells (Feng et al., 2011; Lok et al., 2013). We could detect some residual RAD51 IRIF in the BRCA1-deficient cells, but these were neither reduced by RAD52 depletion, nor by depletion of EXO1, XPF, or ERCC1 (Supplemental figure 3C,D). This indicates that the loss of EXO1 or any of the other SSA factors does not cause lethality in BRCA1-deficient cells by affecting residual HR levels. In contrast, depletion of BRCA2 in the BRCA1-deficient cells resulted in reduced clonogenic outgrowth concomitant with a complete loss of RAD51 IRIF (Figure 6C, Supplemental Figure 3C,D), suggesting that residual HR is essential for survival of BRCA1-deficient cells. Nevertheless, focusing on EXO1, our data indicate that it is its function in SSA, as a back-up pathway for HR, that is essential for the survival of BRCA1-deficient cells.

### BRCA1- and BRCA2-deficient cells display different basal SSA levels

We hypothesized that a reliance on EXO1-mediated SSA could explain the specific lethality of EXO1 loss in BRCA1-versus BRCA2-deficient cells. To test this, we depleted BRCA1 or BRCA2 by RNAi in cells containing our multi-pathway reporter for DSB repair that can quantify repair of a Cas9-induced DSB by SSA and HR simultaneously (van de Kooij et al., 2022). The experiments were performed in EXO1-proficient and -deficient reporter cell lines (Supplemental Figure 3E). As expected, siRNA-mediated depletion of either BRCA1 or BRCA2 led to a full inhibition of HR (Figure 6D left panel, Supplemental figure 3E). In contrast, depletion of these proteins had differential effects on SSA activity (Figure 6D right panel, Supplemental figure 3E). BRCA1 depletion resulted in a decrease in SSA, whereas BRCA2 depletion resulted in an increase in SSA usage, consistent with previous publications (Anantha et al., 2017; Stark et al., 2004; van de Kooij et al., 2022). Importantly, whereas cells lacking both EXO1 and BRCA1 had substantially reduced levels of SSA, concomitant loss of EXO1 did not affect the level of SSA in BRCA2-depleted cells (Figure 6D). These data suggest that BRCA2-deficient cells can still use SSA to overcome DSBs left unrepaired due to defective HR. Indeed, BRCA2-deficient DLD1 cells do not show elevated γH2AX levels upon EXO1 loss (Figure 6E), in contrast to BRCA1-deficient cells (Figure 4E,F). Taken together, the reporter data indicate that EXO1/BRCA1-deficient cells lack HR and have a reduced SSA capacity, while EXO1/BRCA2-deficient cells also lack HR but still have functional SSA.

### BRCA1-mutated tumours show increased usage of SSA

Our data are consistent with a model in which EXO1 loss leads to replication-induced gaps that evolve into DSBs. In BRCA1- or BRCA2-deficient cells these DSBs cannot be repaired by HR. Whereas BRCA2-EXO1-deficient cells can repair these breaks by SSA, allowing survival, BRCA1-EXO1-deficent cells die because SSA-levels are insufficient (Figure 7A). Culminating from this model, *BRCA-* mutated tumours are expected to repair DSBs by EXO1-mediated SSA more frequently than HR-proficient BRCA wildtype tumours. To study this hypothesis, we searched for signatures of SSA usage in genomes of a combined pan cancer cohort, which was divided into HR-proficient, BRCA1-type HR-deficient, or BRCA2-type HR-deficient subgroups, based on algorithmic analysis of genome wide mutational footprints (PCAWG and Hartwig datasets (Martínez-Jiménez et al., 2022; Nguyen et al., 2020)). We defined deletions flanked by homologous sequences of more than 10 bp as genetic ‘scars’ indicative of DSB-repair by SSA. This broad definition will include the majority of SSA repair events while excluding most PolQ-mediated alt-EJ that generally occurs between smaller-sized homology regions (Kelso et al., 2019). In line with our model, the number of genetic SSA scars was higher in BRCA1-type HR-deficient tumour samples compared to HR-proficient tumour samples (Figure 7B). BRCA2-type tumours carried an even higher load of SSA scars than BRCA1-type tumours (Figure 7B), fully consistent with higher levels of SSA in BRCA2-deficient vs BRCA1-deficient cells (Figure 6D). Similar results were obtained when using a more stringent SSA scar definition containing a homology requirement of more than 50 bp (Supplementary figure 3F). Importantly, increased SSA usage correlated with higher *EXO1* expression (Figure 7C). Similarly to what we observed before in breast cancer samples (Figure 2A, B), EXO1 expression was elevated in BRCA1-mutated tumours in this pan cancer cohort as well (Figure 7D). These tumour data are in line with our cellular data showing that BRCA1 deficiency generates a dependency on EXO1-mediated SSA.

**Figure 7.**
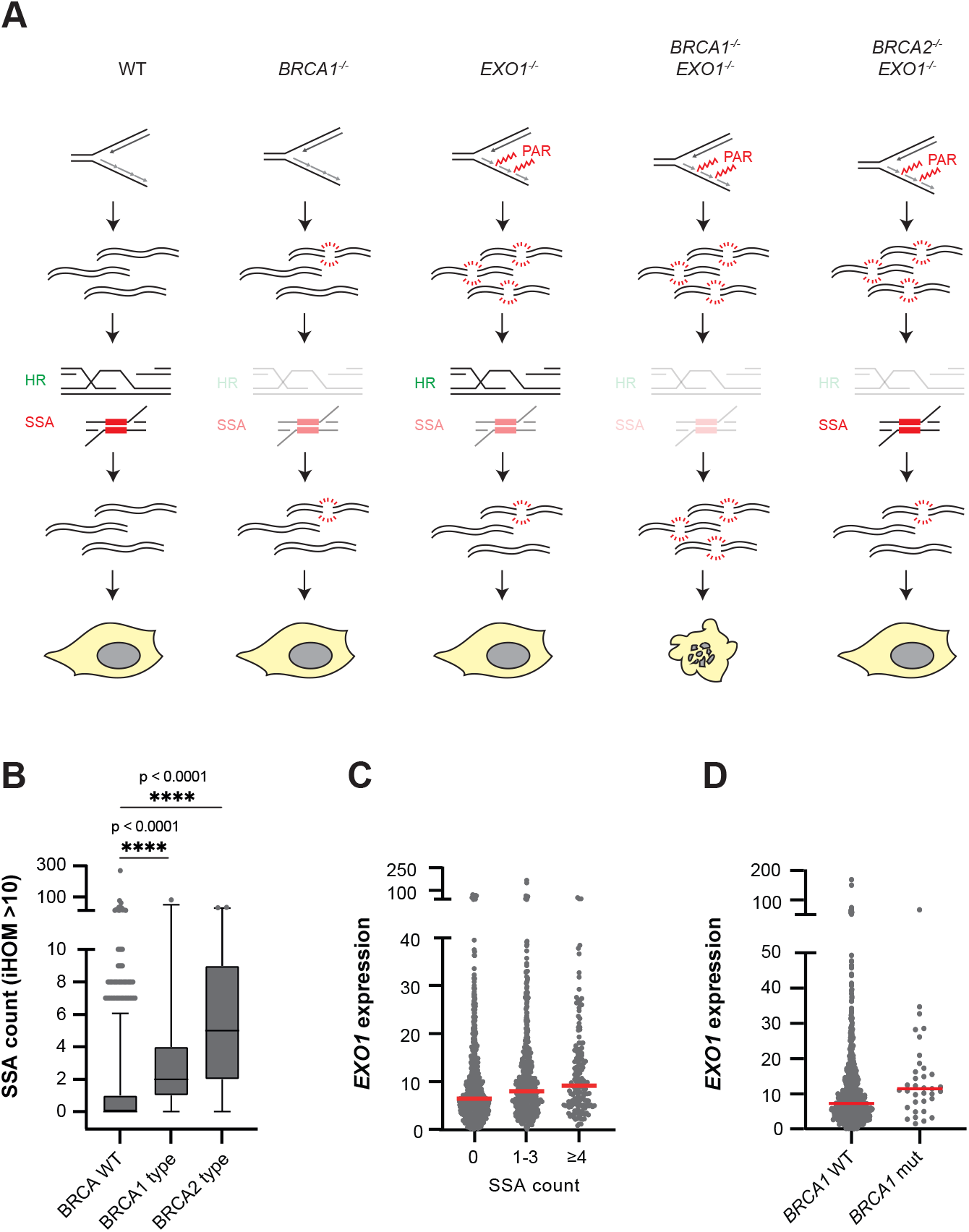
BRCA1-deficient tumours have more SSA-scars than BRCA1-proficient tumours. **(A)** Model of the mechanism causing synthetic lethality between BRCA1-deficiency and EXO1 loss. **(B)** Whole genome sequencing data of pan cancer tumour samples (Martínez-Jiménez *et al.*, 2022) was analysed to quantify the number of genetic scars indicative of DSB-repair by SSA, here defined as deletions flanked by homologous sequences of >10bp. CHORD analysis was used to classify samples as HR-proficient or HR-deficient, either BRCA1-type or BRCA2-type (Nguyen *et al.*, 2020) (****p<0.0001, kolmogornov-smirnov). **(C)** Tumour samples from a pan-cancer cohort were binned based on SSA scar count, and the *EXO1* expression was plotted for each tumour sample. **(D)***EXO1* expression levels in *BRCA1* WT or *BRCA1* mutant pan-cancer tumour samples.

## Discussion

Here, we demonstrate that genetic depletion of EXO1 is severely toxic to BRCA1-deficient cells, identifying a novel target for future therapy development. EXO1 loss in BRCA1-deficient cells increases chromosomal instability which is accompanied by DSB accumulation in S-phase. Our data suggest that these DSBs are left unrepaired in double depleted cells due to an incapacity to perform HR and SSA. These *in vitro* data are substantiated by analysis of tumour samples which indicated that BRCA1-mutated tumours have increased numbers of genetic scars originating from SSA-mediated DSB repair and this correlates with higher levels of EXO1 expression compared to non-mutated tumours.

Unexpectedly, we found that loss of EXO1 is not synthetically lethal with BRCA2 deficiency. Therefore, an HR defect could not be the sole reason for the vulnerability of BRCA1-deficient cells to EXO1 loss. Interestingly, BRCA1-deficient cells show decreased SSA activity, whereas BRCA2-deficient cells show increased SSA activity. The opposing activities of BRCA1 and BRCA2 in SSA have been suggested before and are likely explained by BRCA1, but not BRCA2, driving resection (Densham et al., 2016; Prakash et al., 2015; Stark et al., 2004). We found that cells deficient for BRCA1 and EXO1 have strongly impaired SSA capacity, while EXO1 loss did not reduce the high levels of SSA in BRCA2-deficient cells. These data correlate well with the observed increase of unresolved DSBs in BRCA1-, but not BRCA2-deficient cells upon EXO1 loss. We therefore propose a model in which EXO1 loss results in PAR-decorated post-replicative lesions, that causes further DNA damage, including DSBs. BRCA1-deficient cells that lack EXO1 cannot repair this damage due to a combined HR-deficiency and SSA impairment, resulting in cell death. In contrast, in BRCA2-deficient cells sufficient SSA is retained after EXO1 loss to allow for repair of the DNA damage and survival (Figure 7A). Although further studies are needed to explain why EXO1 loss does not reduce SSA in BRCA2-deficient cells, we hypothesize this relates to differential regulation of resection in BRCA1- and BRCA2-deficient cells. Previously, it has been shown that BRCA1 is required for DNA2 foci formation (Hoa et al., 2015). This might imply that BRCA1-deficient cells cannot stimulate DNA2-mediated resection, thereby becoming more dependent on EXO1 compared to BRCA2-deficient cells that can stimulate resection via both nucleases.

Previous data have shown that BRCA2-deficient cells, as well as BRCA1-deficient cells, depend on RAD52 activity (Feng et al., 2011; Hengel et al., 2016; Huang et al., 2016; Lok et al., 2013). Although RAD52 is a well-known SSA-factor (Rossi et al., 2021), it has also been described to drive BRCA-independent HR (Feng et al., 2011; Lok et al., 2013). In our BRCA1-deficient cells we did not observe any reduction in residual HR upon RAD52 depletion. These data, combined with the observed synthetic lethality of these cells with depletion of RAD52 and other SSA factors, suggest that reduced SSA is at the base of the synthetic lethal interaction between BRCA1 and RAD52 loss. However, such a conclusion also implies that RAD52 is required for SSA in both BRCA1- and BRCA2-deficient cells, unlike EXO1, exemplifying the complexity of the crosstalk between HR and SSA.

Loss of EXO1 strongly increased S-phase PAR levels in our experiments using different cell lines. It has been shown that S-phase PAR is primarily associated with ssDNA gaps resulting from unligated Okazaki fragments (Hanzlikova et al., 2018; Vaitsiankova et al., 2022). Despite the clear increase in PAR-signal in our EXO1-BRCA1-depleted cells, we did not observe long stretches of ssDNA marked by native BrdU staining. This suggests that the DNA lesions represent small gaps or DNA nicks that are below the threshold of BrdU immunofluorescence. In this scenario, one can hypothesize that EXO1 is involved in Okazaki fragment metabolism since it presents flap-endonuclease activity (Keijzers et al., 2015; Lee and Wilson, 1999). Intriguingly, loss of the flap endonuclease FEN1 which is an essential factor in Okazaki fragment maturation, was also found to be synthetically lethal with BRCA-deficiency (Mengwasser et al., 2019). Unlike EXO1, FEN1 is synthetically lethal in both BRCA1- and BRCA2-deficient cells. Although direct evidence for a function of EXO1 in Okazaki fragment maturation in human cells is lacking, experiments in *C. elegans* and *S. cerevisiae* suggest that EXO1 may play an auxiliary role in this process (Kahli et al., 2019; Qiu et al., 1999; van Schendel et al., 2021). While the exact source of the increased PAR-levels upon EXO1 loss remains to be studied, our data suggest that it decorates S-phase DNA lesions that may boost the need for DSB repair pathways since unrepaired ssDNA nicks/gaps can be converted into DSBs when encountered by replication forks (Cortes-Ledesma and Aguilera, 2006; Kuzminov, 2001; Vrtis et al., 2021). In this context, both wildtype and BRCA2-deficient cells can overcome these lesions by HR and SSA, respectively, while BRCA1-deficient cells cannot (Figure 7A).

Similar to SSA, POLQ-mediated alt-EJ has also been shown to be essential in BRCA-deficient cells (Ceccaldi et al., 2015). This suggests that both pathways cannot complement each other to repair unresolved DSBs. Indeed, previous research has shown that both pathways respond to a unique set of DSBs (Kelso et al., 2019). We believe this provides an opportunity to target both pathways simultaneously to improve tumour eradication and reduce risk of resistance to either therapy individually. Next to a role in DSB repair, POLQ has recently been described to be involved in ssDNA gap fill-in (Belan et al., 2022; Mann et al., 2022; Schrempf et al., 2022). However, it is currently unclear how the two different roles of POLQ contribute to its essential role in BRCA-deficient cells. Future research is needed to better understand the nature of the ssDNA gaps that result from inhibition of factors such as POLQ and EXO1, how this relates to their function in DSB repair, and how this drives their synthetic lethal interaction with BRCA deficiency. In conclusion, we show that SSA mediated by EXO1 is essential for BRCA1-deficient cells, but not BRCA2-deficient cells. Therefore, EXO1 is a promising novel target for the treatment of BRCA1-deficient tumours, and the development of inhibitors directed at its nuclease activity should therefore be prioritised.

## Materials and Methods

### Cell lines

RPE1 hTERT *TP53*^-/-^ + FLAG-Cas9 (referred to as WT in this manuscript), RPE1 hTERT *TP53*^-/-^ *BRCA1*^-/-^ + FLAG-Cas9 (referred to as *BRCA1*^-/-^ in this manuscript), RPE1 hTERT *TP53*^-/-^ *BRCA1*^-/-^ *53BP1*^-/-^ + FLAG-Cas9 cells (referred to as *BRCA1*^-/-^ *53BP1*^-/-^ in this manuscript) were previously described (Noordermeer et al., 2018). RPE1 hTERT cells were obtained from ATCC (Manassas, VA, USA). RPE1 hTERT *PAC*^-/-^ *TP53*^-/-^ (referred to as WT in this manuscript) and RPE1 hTERT *PAC*^-/-^ *TP53*^-/-^ *BRCA1*^-/-^ (referred to as *BRCA1*^-/-^ in this manuscript) cells were generated by nucleofection of pLentiCRISPRv2 containing the following sgRNAs: *sgPAC*: ACGCGCGTCGGGCTCGACAT; *sgTP53*:CAGAATGCAAGAAGCCCAGA; sg*BRCA1*: AAGGGTAGCTGTTAGAAGGC. Subsequently, cells were clonally expanded and genotyping was performed by PCR amplification and Sanger sequencing of the targeted locus, followed by TIDE analysis (Brinkman et al., 2014).

The BRCA1 gene in RPE1 hTERT *PAC*^-/-^ *TP53*^-/-^ cells was endogenously C-terminally tagged with *mAID-GFP* as previously described (Sasanuma et al., 2018) (referred to as *BRCA1-mAID* in this manuscript). RPE1 hTERT *TP53*^-/-^ *BRCA1-mAID-GFP EXO1*^-/-^ cells (referred to as *EXO1*^-/-^ in this manuscript) were obtained by nucleofection with pLentiCRISPRv2 containing *sgEXO1* (GCGTGGGATTGGATTAGCAA) and subsequent clonal selection and genotyping as aforementioned. To deplete AID-tagged BRCA1, cells were treated with 500 μM auxin (Sigma Aldrich, Saint Louis, MO, USA; stock solution 35 mg/mL in EtOH) for 48 hours, unless stated otherwise.

HEK 293T cells were obtained from ATCC (Manassas, VA, USA) and HEK 293T DSB-Spectrum V3 cells were previously described (van de Kooij et al., 2022). H1299 pLKO^TetOn^ sh*BRCA2* cells (gift from Dr. Madalena Tarsounas) were previously described (Tacconi et al., 2017). RPE1, HEK 293T and H1299 cells were cultured in Dulbecco’s Modified Eagle’s Medium (DMEM, high glucose, GlutaMAX™ and pyruvate supplemented (ThermoFisher Scientific, Waltham, MA, USA)) + 10% Fetal Calf Serum (FCS) and 1% penicillin + streptomycin (Pen-Strep).

MDA-MB-436 and MDA-MB-436+BRCA1WT cells (gift from Dr. Neil Johnson) and were cultured in Roswell Park Memorial Institute (RPMI) 1640 supplemented with 10% heat-inactivated FCS (Gemini Bio-Products, West Sacramento, CA, USA) and 1% Pen-Strep (ThermoFisher Scientific). Mouse embryonic fibroblasts (MEFs) were previously generated as described in (Callen et al., 2020) and were cultured in DMEM (ThermoFisher Scientific) supplemented with 10% FCS + 1% Pen-Strep. Human DLD-1 WT and *BRCA2*^-/-^ cells (gift from Dr. Stephen Elledge) were maintained in RPMI-1640 Medium (ATCC modification) supplemented with 10% heat-inactivated FCS and 1% Pen-Strep.

All cells were maintained at 37°C, 5% CO2. When BRCA1-deficient, cells were maintained at 37°C, 5% CO2, 3% O2 unless stated otherwise.

### Antibodies

The following primary antibodies were used for western blotting: Mouse α 53BP1 (BD 612522; 1:1,500), Rabbit α BLM (Abcam ab2179; 1:2,000), Mouse α BRCA1 (Merck OP92; 1:1,000), Rabbit α BRCA1 (Millipore 07-434; 1:2,000), Mouse α BRCA2 (Merck OP95; 1:1,000), Mouse α CTIP (Millipore MABE1060; 1:1,000), Rabbit α EXO1 (Abcam ab95068; 1:1,000), Rabbit α EXO1 (Millipore ABE1354; 1:1,000), Mouse α GFP (Sigma #11814460001; 1:1000), Rabbit α MRE11 (1:3,000; de Jager et al., 2001), Mouse α TUBULIN (Sigma T6199; 1:5,000), and Mouse α RAD52 (Santa-Cruz sc-365341; 1:100). The following secondary antibodies were used for western blotting: Goat α Mouse or Goat α Rabbit labelled with respectively IRDye 800 or IRDye 680 (LI-COR; 1:15,000) and HRP-labelled Goat α Mouse and Donkey α Rabbit (ThermoFisher Scientific, 1:5,000).

The following primary antibodies were used for immunofluorescence: Rabbit α BRCA1 (Millipore 07-434; 1:1,000), Mouse α BrdU (Amersham RPN202; 1:800), Rabbit α PAR (Trevigen #4336; 1:500), Rabbit α PAR (Millipore MABE16; 1:250), Rabbit α pRPA32 (Ser4/Ser8) (Bethyl A300-245A; 1:1,000), Rabbit α RAD51 (Bio Academia 70-001; 1:15,000), and Mouse α yH2AX (pSer 139, Millipore 05-636; 1:5,000). The following secondary antibodies were used for immunofluorescence: Goat α Mouse or Goat α Rabbit labelled with either Alexa 488 or Alexa 647 (Invitrogen; 1:1,000).

### Plasmids & cloning

All purification of plasmid DNA or PCR products was done using commercially available kits (Qiagen, Hilden, Germany) according to the manufacturer’s protocol. For the multi-colour competition assays, the sgRNAs targeting the described genes were cloned into the BsmBI-digested lentiviral expression construct pLentiGuide-GFP (Noordermeer et al., 2018). For other experimental purposes, these same sgRNAs were cloned into BsmBI-digested pLentiCRISPRv2 (Sanjana et al., 2014). See Table 1 for used sgRNA sequences. To obtain pCW57.1-BRCA1-3xTy1, full length BRCA1 was PCR-amplified from pCL-MFG-BRCA1 (Addgene #12341 (Ruffner and Verma, 1997)) with a triple Ty1-tag included in the reverse primer and subsenquently cloned into pENTR_1A and transferred to pCW57.1 using gateway cloning. RAD52 was PCR amplified from pEYFP-RAD52 (gift from Jiri Lukas (Bekker-Jensen et al., 2006)) and cloned into pDONR221-eGFP to obtain an eGFP-RAD52 fusion, followed by a gateway LR reaction for transfer into pCW57.1. EXO1 was PCR amplified from pTXB1-EXO1b (Addgene #68267 (El-Shemerly et al., 2005)) and transferred directly into pCW57.1 using Gibson Assembly.

**Table 1.**
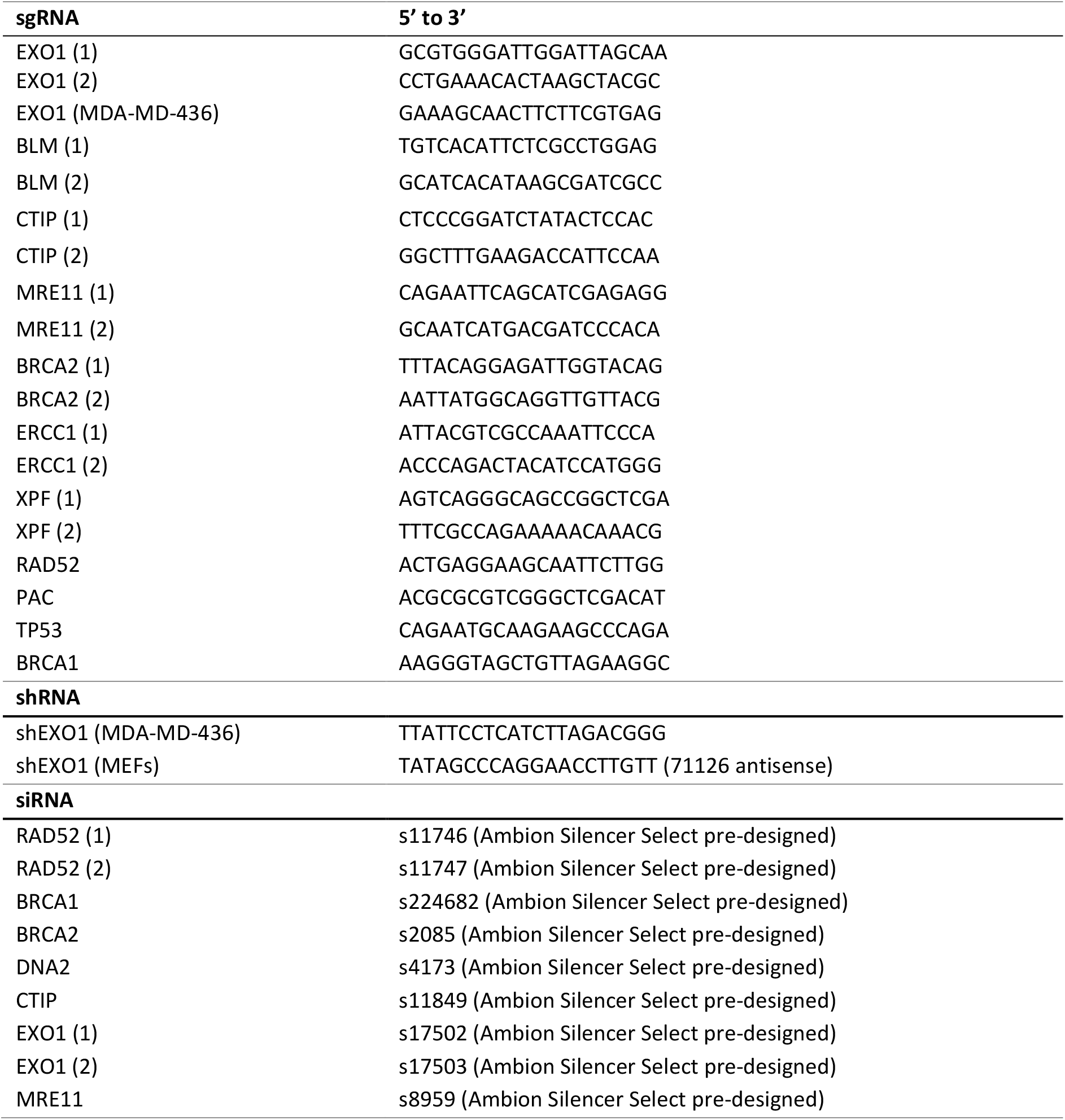
used sgRNAs, shRNAs, and siRNAs throughout this study

For EXO1 knockdown using shRNA, human targeting sh*EXO1* (see Table 1 for sequence, TRC Lentiviral shRNA cloned in pLKO.1, Dharmacon) was used for MDA-MB-436 and mouse targeting sh*EXO1* (see Table 1 for sequence, cloned in pRSITEP-U6Tet-sh-EF1-TetRep-2A-Puro from Cellecta Catalog #: SVSHU6T16-L) was used for MEFs.

### Viral transductions and transfections

Lentivirus was produced in HEK 293T cells by jetPEI transfection (Polyplus Transfection, Illkirch, France) or X-tremeGENE 9 DNA Transfection Reagent (Sigma) of a pLenti, pCW57.1, or shRNA plasmid with third generation packaging vectors pMDLg/pRRE, pRSV-Rev and pMD2.G. Viral supernatants were harvested 48-72 hours post transfection, filtered (0.45 μm filter) and used to transduce cells at an MOI of ~ 1 in the presence of 4 μg/mL or 10 μg/mL polybrene. For RPE1 cells, 10-15 μg/mL puromycin was used for selection, for RPE1 *PAC*^-/-^ cells, 2 μg/mL of puromycin was used, and for MDA-MB4-36 and DLD1 cells 3 μg/mL puromycin was used. Transfections with siRNAs were performed using Lipofectamine RNAiMAX (Invitrogen, Carlsbad, CA, USA) according to the manufacturer’s protocol (see table 1 for used siRNAs). 16 hours post transfection, the medium was refreshed. Transfected cells were used for experiments 48-72 hours after transfection.

### Western Blotting

Cells were lysed in RIPA lysis buffer (1% NP-40, 50 mM Tris-HCl pH 7.5, 150 mM NaCl, 0.1% SDS, 3 mM MgCl2, 0.5% sodium deoxycholate) supplemented with Complete Protease Inhibitor Cocktail (Sigma Aldrich) and 100 U/mL Benzonase (Sigma Aldrich). LDS or SDS sample buffer with DTT was added to the lysates, followed by denaturation at 95°C for 5 minutes. Proteins were separated by SDS-PAGE on 4-12 % gradient gels (ThermoFisher Scientific) or 4-15% Criterion TGX pre-cast midi protein gel (Bio-Rad, Hercules, CA, USA) and transferred to Amersham Protran premium 0.45 μm nitrocellulose membrane (GE Healthcare Life Sciences, Chicago, IL, USA). Membranes were blocked with 5% skimmed milk (Santa Cruz Biotechnology, Dallas, TX, USA) in 1x TBS or Blocking buffer for fluorescent WB (Rockland, Pottstown, PA, USA), and stained with primary and secondary antibodies. After secondary antibody-staining, the membranes were imaged on an Odyssey CLx scanner (LI-COR BioSciences, Milton, UK), followed by image analysis using ImageStudio (LI-COR BioSciences). Alternatively, when HRP-labelled secondary antibodies were used the membranes were treated with the WesternBright ECL HRP Substrate kit (Advansta, San Jose, CA, USA) and imaged on an Amersham Imager 680 (Bioké, Leiden, The Netherlands).

### Multi Colour Competition Assay

RPE1 WT or RPE1 *BRCA1*^-/-^ + FLAG-Cas9 cells were transduced with viral supernatants of pLenti-Guide-GFP containing a sgRNA (see Table 1 for sequences) for the gene of interest (GOI_sgRNA) or pLenti-Guide-mCherry-LacZ_sgRNA. Transduced cells were selected by puromycin treatment for 48h (15 μg/mL for RPE1 WT cells; 10 μg/mL for RPE1 *BRCA1*^-/-^ cells), and subsequently, GFP- and mCherry-positive cells were mixed 1:1. Mixed populations were seeded at 10,000 cells per well in a 12-well plate and imaged every 3 days for a total of 18 days using an ArrayScan Cellomics high content microscope (Thermo Fisher Scientific). HCS Studio Cell Analysis Software (Thermo Fisher Scientific) was used to quantify the number of GFP- and mCherry-positive cells per well. Cells were passaged upon confluency. To assess gene editing efficiencies of the used sgRNAs, whole cell lysates were isolated 5 days post transduction and assessed by Western blotting for the targeted proteins.

### Clonogenics

RPE1 hTERT cells were seeded in 10-cm dishes (RPE1 WT or RPE1 *BRCA1*^-/-^ complemented with BRCA1-WT: 250 cells; RPE1 *BRCA1*^-/-^: 1,000-1,500 cells, RPE1 *BRCA1*^-/-^ *53BP1*^-/-^: 500 cells, RPE1 *BRCA1-mAID*: 500 cells; H1299 pLKO^TetOn^ sh*BRCA2*: 500 cells) and treated as indicated. Medium containing Olaparib (16 nM or 50 nM) (Selleck Chemicals, Planegg, Germany), doxycycline (1 μg/mL) or auxin (500 μM) was refreshed after 7 days. After 14 days, colonies were stained with crystal violet (0.4 % (w/v) crystal violet, 20% methanol) and counted manually.

### Cell Titer Glo

To analyse cell growth and cell viability after EXO1 depletion, MDA-MB-436 and MDA-MB-436+BRCA1WT, DLD1 WT and *BRCA2*^-/-^ cells were transduced with virus containing the *sgEXO1* construction and plated in 6-well plates (10,000 cells per well per sample) after Puromycin selection. Similarly, WT and BRCA1 MEF cells were plated at a density of 10,000 cells per well and doxycycline (1 μg/mL) was added to induce sh*EXO1* expression. The medium was replenished every 3 days and cells were sub cultured when confluent. Cell viability was measured after 12 days using CellTiter-Glo Luminescent Cell Viability Assay (Promega, Madison, WI, USA) following manufacturer’s instructions.

### RNA isolation and qRT–PCR

Total RNA was extracted from cells using Trizol Reagent (Invitrogen), and cDNA was made using SuperScript II Reverse Transcriptase (ThermoFisher), according to manufacturer’s instructions. qPCR was performed using iTaq Universal SyBR Green (Bio-Rad). Samples were run and analysed on a Bio-Rad CFX96 Real-Time PCR detection system. Primer sequences: *mEXO1* forward: 5’ TGTTCGACCCCATCCAAAGG 3’ and *mEXO1* reverse: 5’ GTACTGCCCAGCGTAAGTCA 3’.

### Chromosomal aberrations

Colcemid (0.1 μg/mL) was added to cells at 80% confluency 3 hours prior to harvesting. Cells were harvested by trypsinization and normal medium was added to quench trypsin before centrifugation (5 minutes, 1000 rpm). Supernatant was removed and normal medium was added to create a cell suspension. Freshly made hypotonic solution (1:1 0.4% Na-citrate: 0.4% KCl) was added dropwise to the cell suspension (14:1). Cells were centrifuged 8 minutes, 800 rpm, supernatant was removed partially whereafter 2.5 mL freshly made fixative (3:1 MeOH : Acetic Acid) was added dropwise. Following three washes with the fixative, cells were taken up in fresh fixative. Fixed cells were spread with a Pasteur pipet onto a cleaned microscope slide pre-wet with fixative. Another drop of fixative was added onto the slide with the spread cells at a 45 degrees angle and air-dried overnight. Spreads were stained and mounted using VECTASHIELD^®^ Antifade Mounting Medium with DAPI (Vector Laboratories, Newark, CA, USA). Pictures were made using a Zeiss Axio Imager 2 fluorescent microscope (Zeiss, Oberkochen, Germany) at a 63x zoom. All slides were blinded before quantification of the chromosomal aberrations. At least 40 metaphase spreads per condition per replicate were analysed.

### Micronuclei formation and yH2AX immunofluorescence

Cells were grown in glass bottom 96 well plates (Greiner, Kremsmünster, Austria) until 85% confluency and fixed with 2% (w/v) paraformaldehyde (Sigma Aldrich) in PBS for 20 minutes, washed three times with PBS, and permeabilized with 0.3% Triton X-100 (Sigma Aldrich) in PBS for 20 minutes. Permeabilized cells were subsequently washed three times with PBS and blocked with PBS^+^ (5 g/L BSA (Sigma Aldrich) and 1.5 g/L glycine (Sigma Aldrich) in PBS) for 30 minutes. Blocked cells were incubated 1.5 hours with anti-BRCA1 or anti-yH2AX in PBS^+^, washed 4 times with PBS, and incubated 1 hour with 5 μg/mL Hoechst 34580 (stock 5 mg/mL in H_2_O) (Thermo Fisher) and anti-mouse Alexa 647 in PBS^+^.

To measure yH2AX in EdU positive cells, the cells were incubated 30 minutes at 37°C with EdU (10 μM, stock solution 40 mM) (Invitrogen) before fixation. After staining with the primary and secondary antibody as described above, the cells were fixed with 2% (w/v) paraformaldehyde in PBS for 5 minutes and washed twice with PBS. Subsequently, the fixed cells were incubated with click-it reaction mix (100 mM Tris-HCl pH 8.5, 10 μM Azide-Alexa 647 (Invitrogen), 1 mM CuSO_4_ and 100 mM Ascorbic acid (Sigma Aldrich) added lastly) for 30 min at room temperature.

Cells were washed again 4 times with PBS and analysed using the CellInsight CX7 LZR High content Analysis Platform (Thermo Fisher). For the micronuclei formation, 200 cells per condition per replicate were imaged and the micronuclei were quantified manually. For endogenous yH2AX foci formation 2,000 cells per condition per replicate were imaged and the average nuclear intensity per cell was quantified by the CellInsight CX7 LZR High content Analysis Platform.

### IR-induced foci immunofluorescence

For pRPA IRIF and RAD52 IRIF cells were grown on sterile 13 mm glass coverslips till 85% confluency and fixed 3 hours after irradiation with 10 Gy. Cells were pre-extracted with ice cold nuclear extraction (NuEx) buffer (20 mM Hepes pH 7.5, 20 mM NaCl, 5 mM MgCl2, 1 mM DTT, 0.5% NP-40 (IGEPAL CA-630, Sigma Aldrich), 1x Complete Protease Inhibitor Cocktail (Sigma Aldrich) for 12 minutes at 4°C and directly fixed with 2% (w/v) paraformaldehyde in PBS (20 minutes at room temperature). For RAD51 IRIF, cells were grown on sterile 13 mm glass coverslips till 85% confluency and fixed 3 hours after irradiation with 10 Gy. Cells were fixed and permeabilized with 1% (w/v) paraformaldehyde, 0.5% Triton X-100 in PBS for 20 minutes at room temperature, washed three times with PBS, further fixed and permeabilized with 1% (w/v) paraformaldehyde, 0.3% Triton X-100 in PBS for 20 minutes.

Subsequently, the fixed and permeabilized cells were washed three times with PBS and blocked with PBS^+^ as described above for 30 minutes at room temperature. Blocked cells were incubated 1.5 hours with the primary antibody in PBS^+^, washed 4 times with PBS, and incubated 1 hour with DAPI 0.1 μg/mL (stock 100 μg/mL) and the secondary antibody in PBS^+^. All antibody incubations were performed at room temperature. After washing 4 times with PBS the coverslips were mounted using Aqua-Poly/mount (Polysciences, Warrington, PA, USA). Pictures were made using the Zeiss Axio Imager 2 fluorescent microscope at a 40x zoom. Foci of at least 100 cells per condition per replicate were quantified using the IRIF analysis 3.2 Plugin in ImageJ (Luijsterburg et al., 2016).

### ssDNA analysis

Cells were grown in glass bottom 96 well plates as aforementioned. For BrdU incorporation, the cells were incubated 48 hours at 37°C with BrdU (15 μM, stock solution 30 mM in H2O) (Merck, Darmstadt, Germany). After washing two times with warm PBS the cells were pre-extracted with ice cold NuEx buffer and fixed with 2% (w/v) paraformaldehyde as described previously. For the analysis of PAR levels in EdU-positive RPE1 cells, the cells were incubated 30 minutes at 37°C with PARGi (10 μM, dissolved in DMSO) (MedChemExpress, Monmouth Junction, NJ, USA) and EdU as described before. Thereafter, the cells were pre-extracted with ice cold NuEx buffer supplemented with PARGi and fixed with 2% or 4% (w/v) paraformaldehyde. The fixed and permeabilized cells were blocked and incubated with the appropriate primary, secondary antibody and click-it reaction for EdU as aforementioned. 2,000 cells per condition per replicate were imaged using the CellInsight CX7 LZR High content Analysis Platform. The average nuclear intensity per cell was quantified by the same analysis platform. For the analysis of PAR levels in EdU-positive MDA-MB-436 or DLD1 cells, cells were plated on coverslips pre-treated with 0.1% Gelatin in 6-well plates. Cells were then incubated with EdU for 30 minutes and PARGi (Tocris, Bristol, UK) for 20 minutes before collection. Subsequently, cells were pre-extracted (20 mM Hepes, 50 mM NaCl, 3 mM MgCl2, 0.3 M sucrose, 0.2% Triton X-100) on ice for 5 minutes, fixed with 4% paraformaldehyde (Electron Microscopy Sciences, Hatfield, PA, USA) for 10 minutes and permeabilized using 0.5% Triton X-100 for 10 minutes. The fixed and permeabilized cells were blocked and incubated with the appropriate primary, secondary antibody and click-it reaction for EdU as aforementioned. Images were captured at 40x zoom on a Lionheart LX automated microscope (BioTek, Winooski, VT, USA) or at 63x zoom with an Axio Cam MRC5 attached to an Axio Observer Z1 epifluorescence microscope (Zeiss). Quantification of total nuclear intensity was performed using the Gen5 spot analysis software (BioTek).

### DSB-Spectrum assays

HEK 293T DSB-Spectrum V3 cells were seeded and transfected with siRNAs the next day. A second siRNA transfection was performed 24 hours after the first transfection. 6-8 hours after the second siRNA transfection, 20,000 cells were seeded per well in 96-well plates. 24 hours post-seeding, the cells were transfected in technical duplicate with pX459-Cas9-sgRNA-iRFP construct containing either sgBFP targeting DSB-Spectrum or sgAAVS1. The cells were trypsinised and analysed by flow cytometry 48-96 hours after the DSB-Spectrum targeting transfection. FlowJo software (BD Biosciences, Franklin Lakes, NJ, USA) was used to analyse the acquired flow cytometry data. To select the live, single cell population gating on forward and side-scatter was applied. Subsequently sgAAVS1 or sgBFP targeted cells were selected by gating on iRFP. On this population gating was applied to quantify the frequencies of BFP-/mCherry-, BFP-/mCherry+ and GFP+ cells. The frequency of each fluorescent subpopulation in the sgAAVS1-transfected cells was subtracted from the frequency of that same population in the sgBFP-transfected cells. The resulting background-corrected frequencies were normalized to the *siCTRL* transfected cells.

### Clinical data analyses

Correlations between *EXO1* and *BLM* expression and *BRCA1* mutations was studied in two different previously published breast cancer cohorts. The first cohort contained 247 patients diagnosed with triple-negative breast cancer of which 22 tumours harboured biallelic *BRCA1* mutations and 59 tumours displayed *BRCA1* promotor hypermethylation (Staaf et al., 2019). The second contained 342 patients diagnosed with breast cancer of which 15 tumours harboured deleterious BRCA1 mutations (Nik-Zainal et al., 2016).

The Xena functional genomic platform (Goldman et al., 2020) was used to extract both genome wide gene expression levels (FPKM) as well as the HRD-score (Knijnenburg et al., 2018) for all samples in the TCGA breast cancer cohort. Tumour samples were excluded from analysis in case HRD-score and/or gene expression data were absent. If multiple tumour samples from a single patient were sequenced, as was the case for six patients, the expression values were averaged. Next, for each individual gene, the Pearson correlation coefficient between gene expression level and HRD score was determined.

A unified whole genome mutational dataset of the Hartwig Medical Foundation (Hartwig) and the Pan-Cancer Analysis of Whole Genomes (PCAWG) cohort was used to search for SSA scars (Martínez-Jiménez et al., 2022). HR-status was classified according to genome analysis using the CHORD algorithm as previously described (Nguyen et al., 2020). The structural variants (SV) of this dataset were called with the genomic rearrangement toolkit LINX (10.1016/j.xgen.2022.100112). This tool integrates copy number profiles and the SV calls from PURPLE (0.1038/s41586-019-1689-y) and GRIDSS (10.1186/s13059-021-02423-x) that enables the characterization and annotation of simple and complex genomic rearrangements. Specifically, LINX chains one or more SVs and classifies these SV clusters into various event types (‘ResolvedType’). We defined deletions and duplications as clusters with a ResolvedType of ‘DEL’ or ‘DUP’ whose start and end breakpoints were located on the same chromosome (*i.e.* intrachromosomal). Duplication or deletion events that were part of a complex SV event were excluded from this analysis as these are not induced by a continuous SV-related mutation process. Subsequently, each duplication or deletion event was annotated with its homology sequence length at the break junction using IHOMPOS from PURPLE. This SV mutation feature represents the imperfect homology length of a break junction left and right from the breakend and is calculated by performing local Smith-Waterman alignment of the breakpoint sequence to the reference sequence up to 300bp either side of the break junction. The total homology length of a break-end was calculated as the sum of the left and right IHOMPOS sequence length around the breakend. Lastly, the logics of the PCAWG SV reference paper (Li et al., 2020) were used to classify SV scars. Duplication events were subsequently excluded from analysis, and all deletions bearing an (imperfect) homology sequence length longer than 10 or 50 base pairs were annotated as SSA mutation scars. The cut-off of 10 bps was chosen to rule out inclusion of NHEJ or alt-EJ scars, which are generally dependent on shorter stretches of homology.

## Acknowledgements

We would like to thank Hiroyuki Sasanuma (Kyoto University, Japan) for sharing plasmids to generate BRCA1-mAID cells and Johan Staaf (Lund University, Sweden) for sharing expression data on triplenegative breast cancer cases. We also would like to thank Dr. Madalena Tarsounas (University of Oxford, UK) for sharing the H1299 pLKO^TetOn^ sh*BRCA2* cells, Dr. Neil Johnson for sharing the MDA-MB-436 and MDA-MB-436+BRCA1WT cells, and Dr. Stephen Elledge for sharing the Human DLD-1 WT and BRCA2-deficient cells. This work was funded by grants from the Dutch Research Council (NWO, VIDI grant #192.039 to SMN), Oncode Institute (SMN), Leiden University Medical Center (LUMC, MSCA-IF Seal of Excellence to BvdK) and the Dutch Cancer Foundation (KWF, research grant #13472 to SMN and HvA; personal grant #BUIT 2015-7546 to BvdK). We thank Andrew Blackford for helpful discussions on this project. This publication and the underlying study have been made possible partly based on data that Hartwig Medical Foundation has made available to the study through the Hartwig Medical Database.

## Datasets

The Hartwig data was provided under data transfer agreement DR-247 from the Hartwig Medical Foundation. The PCAWG-US was approved by National Institutes of Health (NIH) for the dataset General Research Use in The Cancer Genome Atlas (TCGA) on 25 February 2021 under application number 100344-3. Access to the non-US PCAWG samples was granted via the Data Access Compliance Office (DACO) Application Number DACO-1050905 on 6 October 2017.

## Author contributions

The project was initiated by BvdK and SMN. Experiments were performed and analyzed by BvdK, AS, RSP, VG, MSMA, DK, TJW, EC, and SMN and conceptualized by BvdK, AS, RSP, AN, HvA, and SMN. *In silico* analyses of tumour material was performed and interpreted by BvdK, AvH, JB, HM, EC, and SMN. The manuscript was written by AS and SMN with help from BvdK, RSP, AN, and HvA and input from all other co-authors.

