## Supplemental Figures for "EXO1-mediated DNA repair by single-strand annealing is essential for BRCA1-deficient cells"

### Supplemental Figure 1

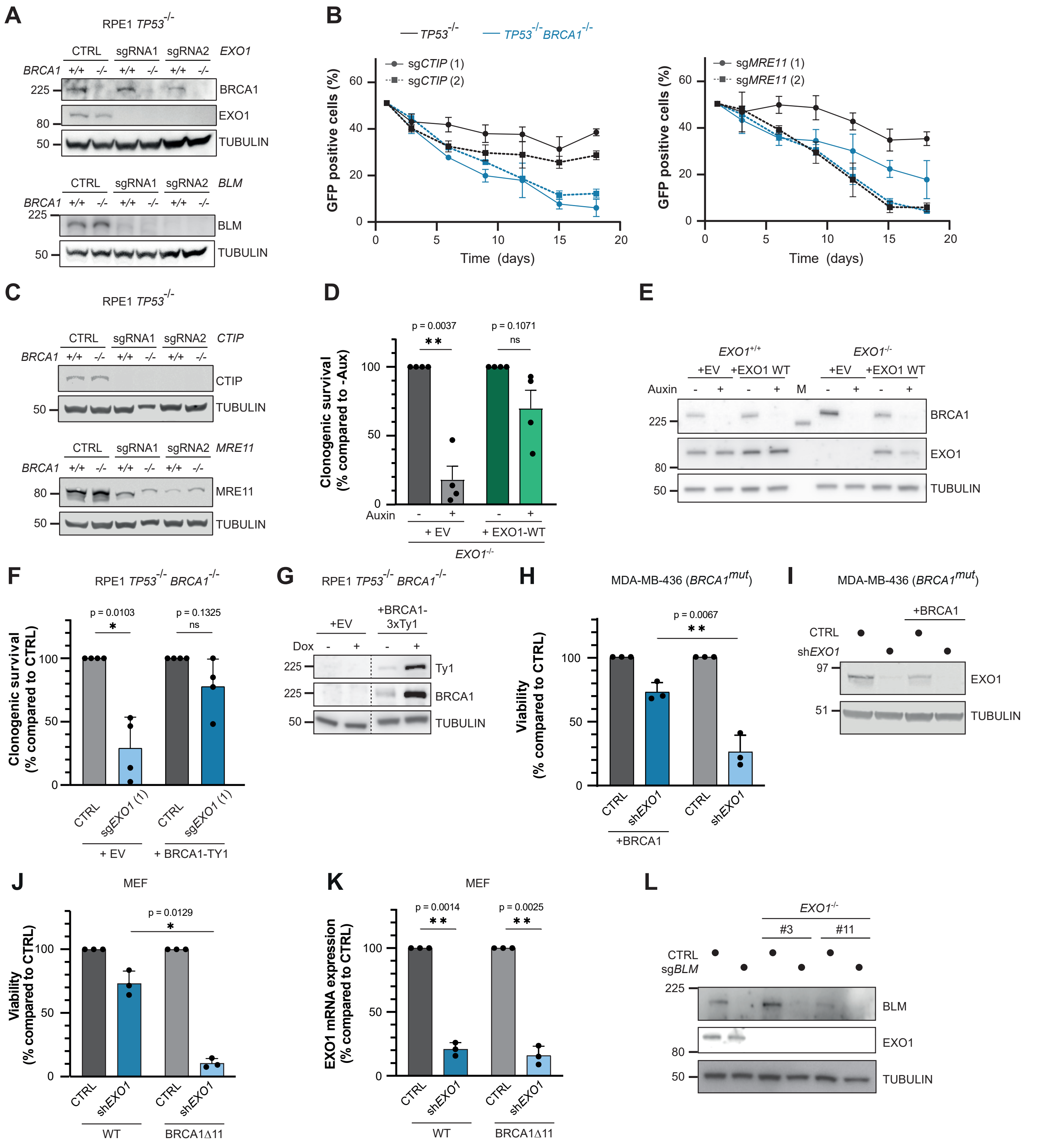


**Supplemental figure 1. Synthetic lethality of EXO1 loss in BRCA1-depleted cells is rescued by re-expression of either factor.** **(A)** Western blot of lysates from the cells analysed in figure 1B. **(B)** As in figure 1B (n=4, mean±SEM). **(C)** Western blot of lysates from the cells analysed in supplemental figure 1B. **(D)** *EXO1* cDNA, or an empty vector (EV) control, were lentivirally transduced into RPE1 hTERT *TP53*$^{-/-}$ *BRCA1-mAID-GFP* cell lines that were either *EXO1*$^{+/+}$ or *EXO1*$^{-/-}$. A clonogenic survival assay was performed in the presence or absence of 500 μM auxin (n=4, mean±SEM, **p<0.01, paired t-test). **(E)** BRCA1 and EXO1 expression in the cells analysed in supplemental figure 1D was determined by western blotting. M=protein ladder **(F)** RPE1 hTERT *TP53*$^{-/-}$ *BRCA1*$^{-/-}$ cells, virally reconstituted with a doxycycline-inducible *BRCA1-Ty1* cDNA, or an empty vector (EV) control, were lentivirally transduced with *Cas9* cDNA and either an AAVS1-targeting (CTRL) or *EXO1*-targeting sgRNA. Next, cells were treated with doxycycline, followed by a clonogenic survival assay (n=4, mean+SD, *p<0.05, paired t-test). **(G)** Western blot analysis of BRCA1 expression in the cell lines used in supplemental figure 1F. Samples were run on the same gel, dashed line indicates removal of non-relevant lanes post-imaging. **(H)** *BRCA1*-mutated MDA-MB-436 cells, either WT or reconstituted with BRCA1, were infected with empty vector (CTRL) or *EXO1*-targeting shRNA and viability was measured using CellTiter-Glo. (n=3, mean+SD, paired t-test). **(I)** Western blot of lysates from the cells analysed in supplemental figure 1H. **(J)** Mouse Embryonic Fibroblasts (MEFs) with a homozygous exon 11 deletion in the *BRCA1* gene (Δ11) were infected with control or *EXO1*-targeting shRNA, followed by a viability assay (n=3, mean+SD, *p<0.05, paired t-test). **(K)** *EXO1* expression was determined by qPCR analysis in the cells analysed in supplemental figure 1J (n=3, mean+SD, **p<0.01, paired t-test). **(L)** Western blot of lysates from the cells analysed in figure 1G.

#### Supplemental Figure 2

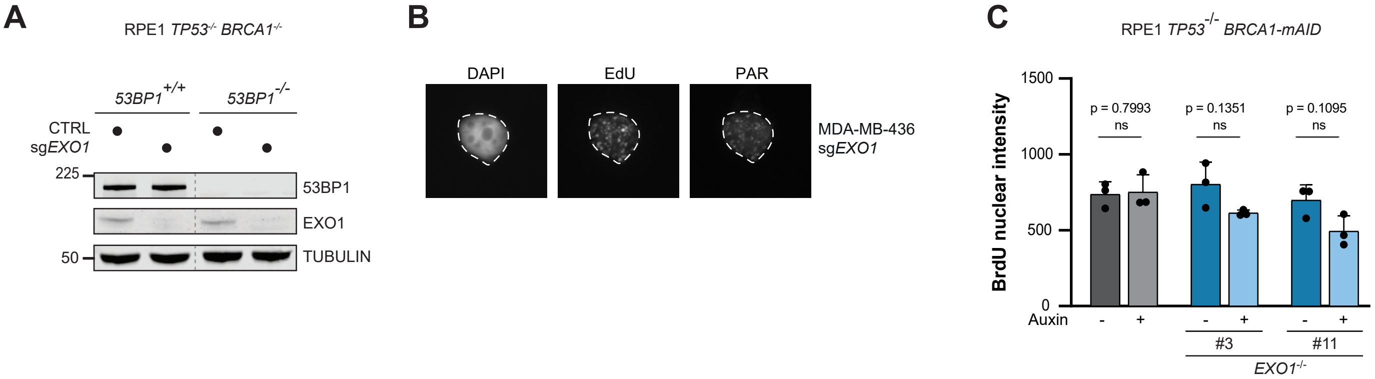


**Supplemental figure 2. Increased S-phase PAR in *EXO1*$^{-/-}$ cells. (A)** Western blot of lysates from the cells analysed in figure 5A. Samples were run on the same gel, dashed line indicates removal of non-relevant lanes post-imaging. **(B)** Repre-representative microscopy images (figure 5C) of the co-localisation between PAR and EdU. **(C)** RPE1 hTERT *TP53*$^{-/-}$ *BRCA1-mAID-GFP* cell lines, either *EXO1*$^{+/+}$ or *EXO1*$^{-/-}$, were treated with 500 μM auxin or left untreated. Cells were incubated with BrdU for 48 hours, followed by native anti-BrdU staining and IF microscopy to quantify average nuclear BrdU intensity. At least 500 cells per condition per replicate were analysed (n=3, mean+SD, paired t-test).

#### Supplemental Figure 3

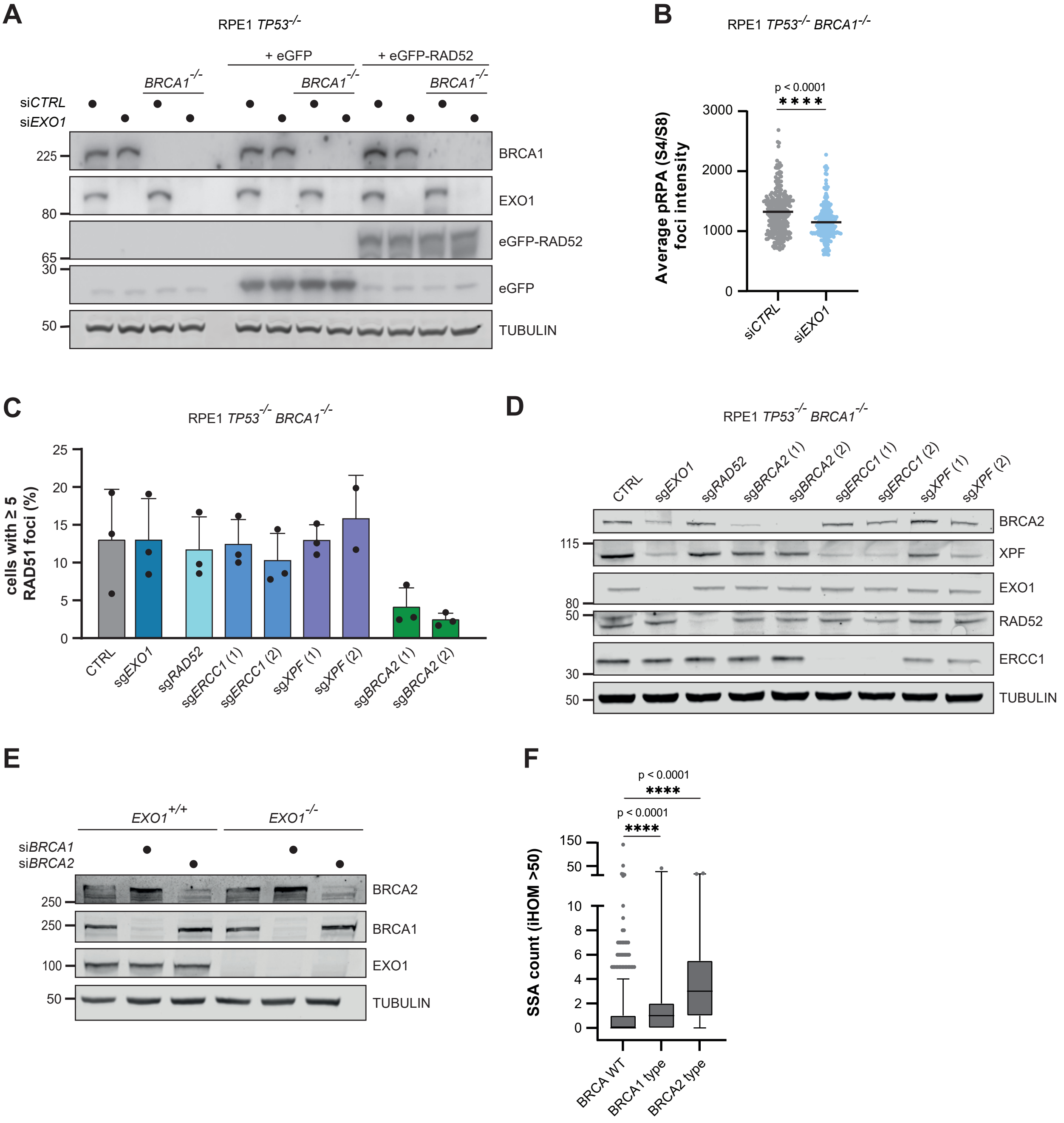


**Supplemental figure 3. Loss of EXO1 affects resection, not HR. (A)** Western blot of lysates from the cells analysed in figure 6B and supplemental figure 3B. **(B)** RPE1 hTERT *TP53*$^{-/-}$ *BRCA1*$^{-/-}$ cell lines were transfected with a control (si*CTRL*) or *EXO1*-targeting siRNA. Next, cells were treated with 10 Gy of IR, followed by IF microscopy to quantify phosphorylated (S4/8) RPA foci. At least 100 cells per condition per replicate were analysed (n=3, mean, ****p<0.01, kolmogornov-smirnov). **(C)** Cell lines depicted in figure 6C were treated with IR, followed by IF microscopy to quantify RAD51 foci (n=3, mean±SD). **(D)** Western blot of lysates from the cells analysed in figure 6C and supplemental figure 3C. **(E)** Western blot of lysates from the cells analysed in figure 6D. **(F)** As in figure 7B, but now SSA-scars are defined as deletions flanked by homologous sequences of >50bp (****p<0.0001, kolmogornov-smirnov).
